# Clinical and Genomic Characterization of Recalcitrant Enterococcal Bacteremia: A Multicenter Prospective Cohort Study (VENOUS)

**DOI:** 10.1101/2025.04.01.645485

**Authors:** Shelby R. Simar, Truc T. Tran, Kirsten B. Rydell, Rachel L. Atterstrom, Pranoti V. Sahasrabhojane, An Q. Dinh, Marissa G. Schettino, Haley S. Slanis, Alex E. Deyanov, Andie M. DeTranaltes, Dierdre B. Axell-House, William R. Miller, Jose M. Munita, David Tobys, Harald Seifert, Lena Biehl, Marcus Zervos, Geehan Suleyman, Jagjeet Kaur, Victoria Warzocha, Renzo O. Cifuentes, Lilian M. Abbo, Luis Shimose, Catherine Liu, Katherine Nguyen, Ashleigh Miller, Samuel A. Shelburne, Blake M. Hanson, Cesar A. Arias

**Affiliations:** Center for Infectious Diseases, UTHealth-Houston School of Public Health, Houston, TX, USA; Division of Infectious Diseases, Houston Methodist Hospital, Houston, TX, USA; Center for Infectious Diseases, Houston Methodist Research Institute, Houston, TX, USA; Department of Infectious Diseases, Division of Internal Medicine, The University of Texas MD Anderson Cancer Center, Houston, TX, USA; Genomics and Resistant Microbes Group, Facultad de Medicina Clínica Alemana de Santiago, Universidad del Desarrollo, Santiago, Chile; Institute for Medical Microbiology, Immunology and Hygiene, Faculty of Medicine and University Hospital, Cologne, University of Cologne, Cologne, Germany; German Center for Infection Research (DZIF), Partner Site Bonn-Cologne, Cologne, Germany; Institute of Translational Research, Cologne Excellence Cluster on Cellular Stress Responses in lAging-Associated Diseases (CECAD), Faculty of Medicine and University Hospital Cologne, University of Cologne, Cologne, Germany; Department of Internal Medicine, Faculty of Medicine and University Hospital of Cologne, University of Cologne, 50924, Cologne, Germany; Department of Internal Medicine, Division of Infectious Diseases, Henry Ford Hospital, Detroit, MI, USA; Jackson Health System, Miami Transplant Institute, Miami, FL, USA; Division of Infectious Disease, Department of Medicine, University of Mississippi Medical Center, Jackson, MS, USA; Department of Medicine, Division of Allergy and Infectious Diseases, School of Medicine, University of Washington, Seattle, WA, USA; Vaccine and Infectious Disease Division, Fred Hutchinson Cancer Center, Seattle, WA, USA; Department of Medicine, Weill Cornell Medical College, New York, NY, USA

**Keywords:** *Enterococcus*, bacteremia, persistence, genomic adaptation

## Abstract

**Background:** Patients with recalcitrant enterococcal bloodstream infections are at greater risk of adverse outcomes. We identified patients in the 2016-2022 Vancomycin-Resistant Enterococcal Bacteremia Outcomes Study (VENOUS) cohort experiencing recalcitrant bloodstream infections for further clinical and genomic characterization.

**Methods:** Bacteremia episodes were considered “persistent” if there was a lack of clearance on day four while receiving ≥ 48 hours of active therapy and recurrent if there was clearance during hospitalization with a subsequent positive culture (collectively, “recalcitrant” bacteremia). A matched comparison group of non-recalcitrant bacteremia patients was chosen in a 2:1 control:case ratio. Isolates were subjected to short- and long-read whole-genome sequencing. Hybrid assemblies were created using a custom pipeline. *Findings.* A total of 46 recalcitrant infections from 41 patients were identified. Patients with persistent bacteremia were more often admitted to the ICU upon admission relative to controls. *E. faecalis* strains causing persistent infections had a significantly higher proportion of genes associated with carbohydrate utilization relative to controls. Representation of functional groups associated with mutated genes was disparate between *E. faecium* and *E. faecalis* index and persistent isolates, suggesting species-specific adaptation.

**Discussion:** Enterococcal isolates causing recalcitrant bacteremia were genomically diverse, indicating that strain-specific signatures are not drivers of persistence. However, comparisons of index vs. persistent isolates revealed that *E. faecium* may be genetically pre-adapted to cause persistent infection, and site-specific structural variation during infection suggests the role of differential gene expression in adaptation and persistence. This data lays groundwork for future studies to define signatures of enterococcal adaptation during bacteremia.

## INTRODUCTION

Persistent enterococcal bacteremia is common in clinical practice, despite the administration of seemingly appropriate anti-enterococcal antibiotic therapy.^1^ Importantly, recent studies have shown that failure to clear enterococcal BSI is associated with a higher in-hospital mortality.^2,3^

Persistent enterococcal bacteremia has been attributed to “translocation” from the gastrointestinal tract, particularly in patients with cancer and neutropenia and those subjected to liver transplants.^4^ Additionally, persistent positive blood cultures with enterococci are often associated with infected vascular devices or endovascular infections with no source control.^5^ Once in the bloodstream, these organisms are subjected to niche-specific selective pressures, which may select for hardier and more adaptable strains that allow for persistence in this new niche.^6^

Studies that comprehensively describe the population genomics and clinical factors associated with persistent enterococcal bacteremia are scarce and are generally limited to small case studies of patients at a single institution. Here, we aimed to comprehensively characterize persistent and recurrent enterococcal bacteremia in a multicenter cohort (Vancomycin-Resistant Enterococcal BSI Outcomes Study; VENOUS).

## METHODS

### Source population and eligibility criteria

VENOUS is a global, prospective, observational study of adults (≥18 years of age) admitted to participating institutions from 2016-2023 with at least one blood culture positive for *Enterococcus faecalis or Enterococcus faecium*, along with one follow-up blood culture taken within seven days following the initial positive culture. Clearance of bacteremia was defined as two consecutive negative cultures, or one negative culture and no positive cultures in the following four days. Patients were considered as having persistent bacteremia if there was a lack of clearance on day four while receiving at least 48 hours of active anti-enterococcal therapy (previously defined as “microbiological failure” in VENOUS) ^2^. Those with recurrent bacteremia either had clearance of bacteremia during hospitalization, with a second positive culture also during this period, or two negative cultures at least one day apart, followed by at least one positive culture. Persistent and recurrent episodes were collectively referred to as “recalcitrant” bacteremia. Index isolates and isolates from the positive culture at the time of persistence or recurrence were used for genomic analyses. A comparison cohort of patients with total clearance of bacteremia per clinical documentation was also selected in a 2:1 ratio of non-recalcitrant:recalcitrant bacteremia (**Figure 1**). Groups were balanced with respect to hospital of admission, timing of the index positive culture, and enterococcal species (*E. faecalis* or *E. faecium*).

**Figure 1.**
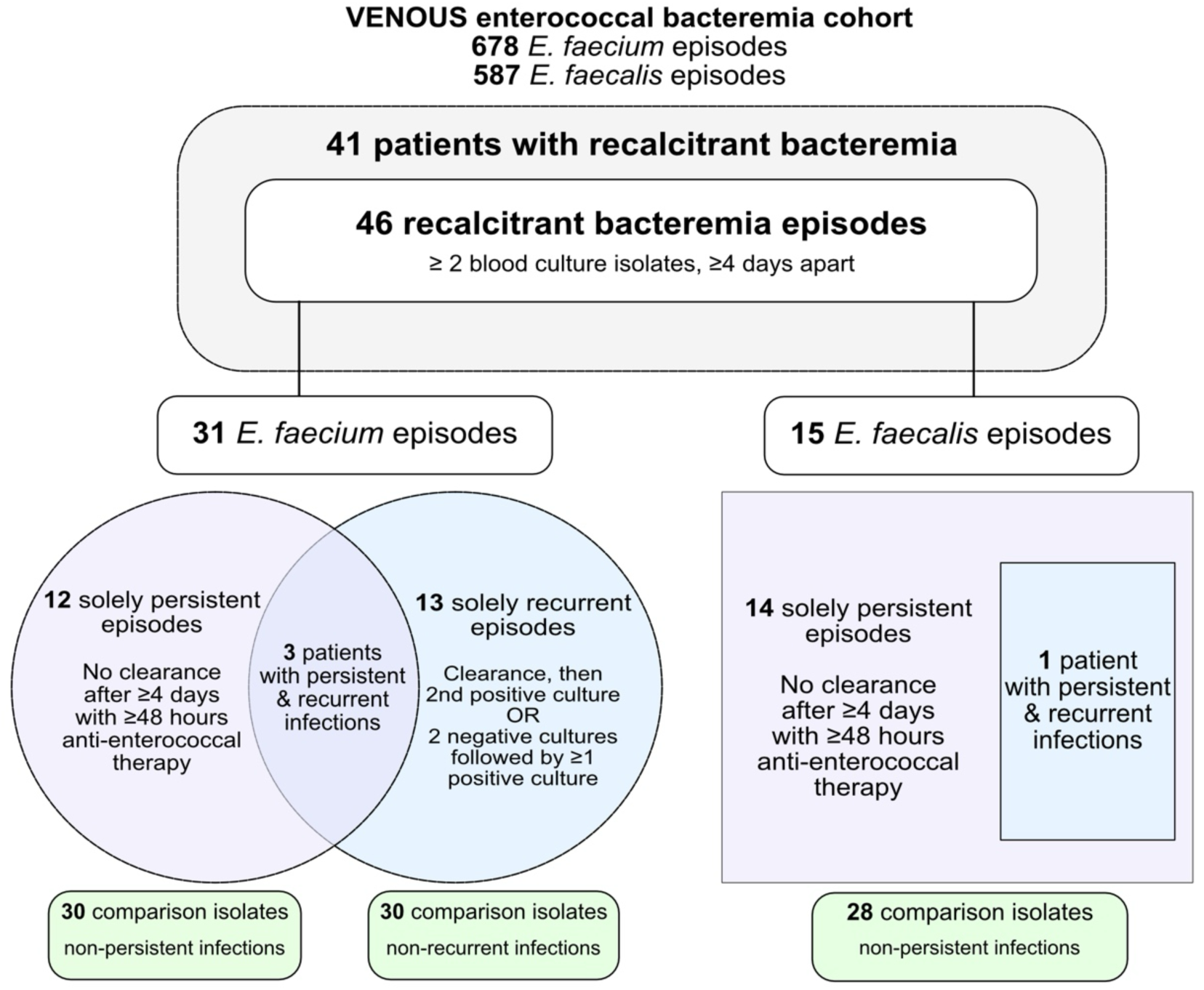
Flow diagram of the study and isolate selection. This figure depicts the identification and selection criteria of recalcitrant bacteremia episodes from the VENOUS source population, along with the number of infections defined as persistent, recurrent or both, for each enterococcal species.

**Figure 2.**
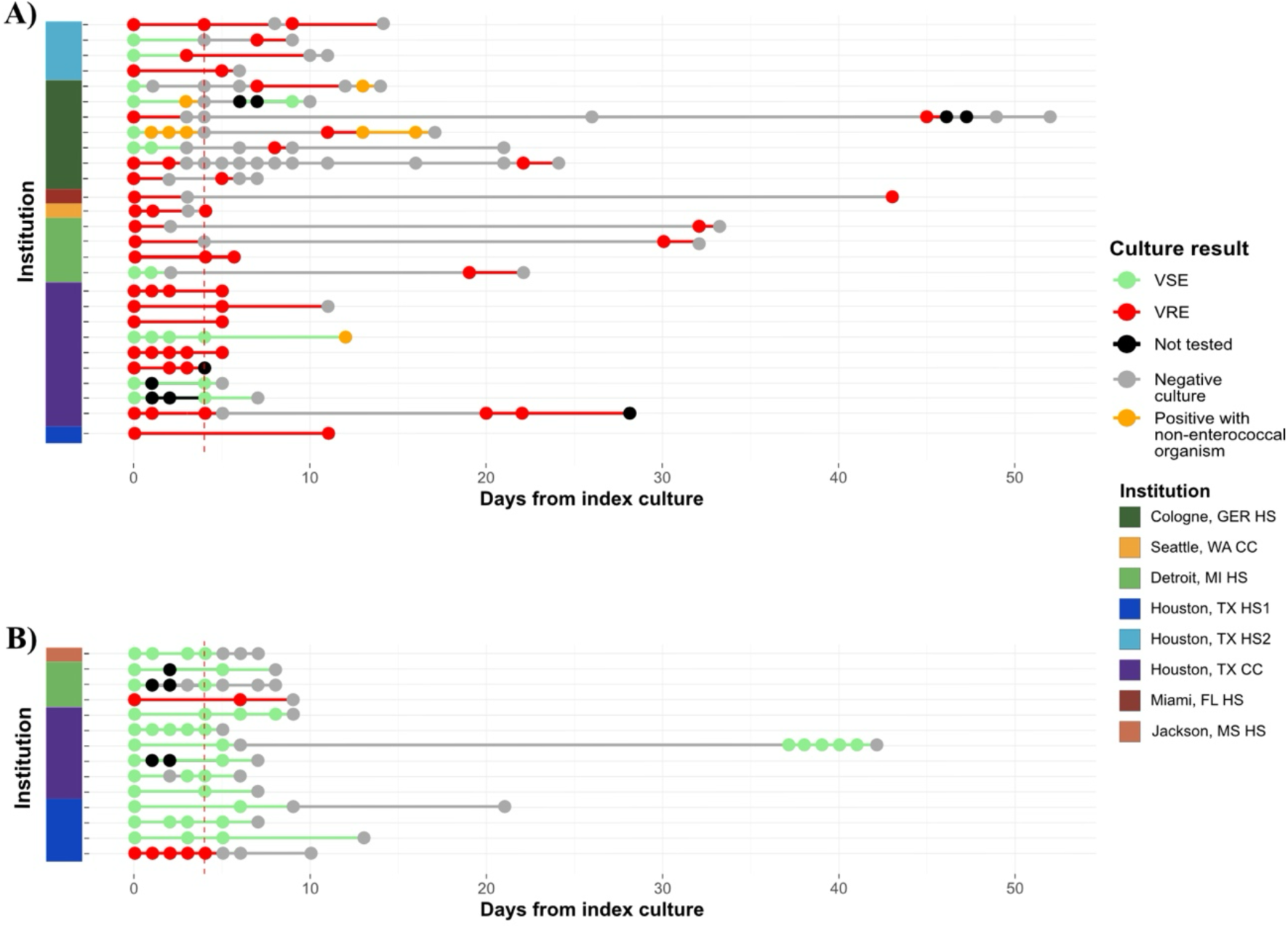
Timeline of recalcitrant bacteremia episodes relative to date of hospital admission. (A) *E. faecium.* (B) *E. faecalis.* Each horizontal line represents a single patient’s hospital course from index culture to the day of the persistent or recurrent culture, and clearance of the bacteremia if documented. The vertical dotted red line indicates day four following collection of the index blood culture, which is designated as time 0 on this chart. VSE = vancomycin-susceptible enterococci; VRE = vancomycin-resistant enterococci; HS = hospital system; CC = cancer center

### Clinical data collection

Clinical, microbiological, and demographic data were collected from electronic medical records using REDCap (Vanderbilt University). Pitt bacteremia score and Charlson Comorbidity Index were calculated for each patient based on clinical data values.^7,8^ Details on empiric and definitive therapy definitions can be found in **Supplementary Material**.

### Population structure and phylogenetic analyses

Isolate processing, storage, sequencing, assembly, and genomic analyses are described in **Supplementary Material**. The pan-genome of final hybrid assemblies was reconstructed for *E. faecium* and *E. faecalis* separately using Roary-v3.11.2.^9^ The core genome alignment file produced from Roary was used as input for separate midpoint-rooted maximum-likelihood phylogenetic tree creation using RAxML-v8.2.12^10^ with 100 bootstrap iterations. Trees were visualized using iTOL.^11^

### Statistical analysis of clinical and genomic data

Comparisons between persistent or recurrent bacteremia patients and their respective comparison cohorts, as well as comparisons between persistent infection pairs (the index isolate and the “persistent time point” isolate collected at day ≥4 of bacteremia) were performed separately for *E. faecium* and *E. faecalis* using R-v4.1.2.^12^ Details on specific analyses are included in **Supplementary Material**, while the results of these analyses can be found in **Supplementary Data 1, 2**. The study was approved by the local institutional review board of participating institutions, which waived the requirement for written or verbal consent from patients based on the observational nature of the study.

## RESULTS

Amongst 1,265 unique patients with enterococcal bacteremia (678 *E. faecium*; 587 *E. faecalis*), we identified 41 patients with recalcitrant bacteremia episodes from eight study sites (**Figures 1, 2**). Of these, there were 29 solely persistent bacteremia episodes, 13 exclusively recurrent bacteremia episodes, and three persistent followed by recurrent bacteremia episodes (included in both persistent and recurrent clinical data analyses). The median length of time between index blood culture and persistent culture was four days (IQR 4; 5) with a nearly even split between patients with *E. faecium* and *E. faecalis* (51.7% and 48.3%, respectively). Recurrent infections had a median time between index and recurrent blood cultures of 26 days (IQR 7; 34) and were predominately caused by *E. faecium* (93.8%) (**Tables 1, 2**). A total of 58 patients without persistent or recurrent bacteremia served as a control group for the persistent bacteremia cohort, and 30 patients were included as controls for the recurrent bacteremia cohort. Of note, matching by institution for one study site was not possible due to lower patient enrollment at the time of analysis.

**Table 1.** Clinical characteristics of patients with persistent enterococcal bacteremia and comparison patients.

| Variables | Persistent<br>(n=29) | Comparison<br>(n=58) | Total Population<br>(n= 87) | Univariable<br>p-value | GLMM p-value |
| --- | --- | --- | --- | --- | --- |
| Institution |  |  |  | N/A;<br>matching<br>factor |  |
| Seattle, WA CC | 1 (3.4) | 2 (3.4) | 3 (3.4) |  |  |
| Houston, TX CC | 15 (51.7) | 30 (51.7) | 45 (51.7) |  |  |
| Detroit, MI HS | 4 (13.8) | 8 (13.8) | 12 (13.8) |  |  |
| Houston, TX HS 1 | 5 (17.2) | 10 (17.2) | 15 (17.2) |  |  |
| Houston, TX HS 2 | 3 (10.3) | 6 (10.3) | 9 (10.3) |  |  |
| Jackson, MS HS | 1 (3.4) | 2 (3.4) | 3 (3.4) |  |  |
| Demographics |  |  |  |  |  |
| *Age, years* |  |  |  |  |  |
| ≤40 | 6 (20.7) | 13 (22.4) | 19 (21.8) | 1 |  |
| 41-50 | 2 (6.9) | 5 (8.6) | 7 (8.0) | 1 |  |
| 51-60 | 4 (13.8) | 11 (19.0) | 15 (17.2) | 0.76 |  |
| 61-70 | 13 (44.8) | 20 (34.5) | 33 (37.9) | 0.48 |  |
| 71-80 | 4 (13.8) | 9 (15.5) | 13 (14.9) | 1 |  |
| *Race* |  |  |  |  |  |
| Asian | 2 (6.9) | 2 (3.4) | 4 (4.6) | 0.60 |  |
| Black or African-American | 3 (10.3) | 13 (22.4) | 16 (18.4) | 0.28 |  |
| Caucasian/White | 21 (72.4) | 37 (63.8) | 58 (66.7) | 0.57 |  |
| Unknown | 3 (10.3) | 6 (10.3) | 9 (10.3) | 1 |  |
| *Ethnicity* |  |  |  |  |  |
| Hispanic | 6 (20.7) | 5 (8.6) | 11 (12.6) | 0.21 |  |
| Not Hispanic | 20 (69.0) | 47 (81.0) | 67 (77.0) | 0.32 |  |
| Unknown | 3 (10.3) | 6 (10.3) | 9 (10.3) | 1 |  |
| Sex, male | 21 (72.4) | 27 (46.6) | 48 (55.2) | **0.04** | 0.48 |
| Current admission |  |  |  |  |  |
| Intensive care unit admission | 14 (48.3) | 14 (24.1) | 28 (32.2) | **0.04** | 0.70 |
| Reason of admission-medical/non-surgical | 27 (93.1) | 56 (96.6) | 83 (95.4) | 0.86 |  |
| Length of hospitalization, days, median (range) | 37 (10-354) | 21.5 (3-101) | 25 (3-354) | **<0.001** | **<0.001** |
| Medical history |  |  |  |  |  |
| *Baseline comorbidities* |  |  |  |  |  |
| Heart/cardiovascular disease | 2 (6.9) | 3 (5.2) | 5 (5.7) | 1 |  |
| Diabetes mellitus | 12 (41.1) | 18 (31.0) | 30 (34.5) | 0.47 |  |
| Chronic obstructive pulmonary disease | 3 (10.3) | 7 (12.1) | 10 (11.5) | 1 |  |
| Chronic kidney disease | 8 (27.6) | 8 (13.8) | 16 (18.4) | 0.20 |  |
| Liver disease | 4 (13.8) | 6 (10.3) | 10 (11.5) | 0.91 |  |
| Solid malignancy | 8 (27.6) | 13 (22.4) | 21 (24.1) | 0.79 |  |
| Hematological malignancy | 13 (44.8) | 23 (39.7) | 36 (41.4) | 0.82 |  |
| Charlson Comorbidity Index, median (range) | 8 (4-13) | 7 (2-12) | 8 (2-13) | 0.50 |  |
| Solid organ transplant | 3 (10.3) | 1 (1.7) | 4 (4.6) | 0.21 |  |
| Bone marrow transplant | 6 (20.7) | 3 (13.8) | 14 (16.1) | 0.61 |  |
| Immunosuppressive therapy | 20 (69.0) | 32 (55.2) | 52 (59.8) | 0.25 |  |
| Cardiac device and cardiac valve | 4 (13.8) | 4 (6.9) | 8 (9.2) | 0.43 |  |
| Hemodialysis prior to admission | 1 (3.4) | 1 (1.7) | 2 (2.3) | 1 |  |
| Previous hospitalization | 24 (82.8) | 43 (74.1) | 67 (77.0) | 0.53 |  |
| within 365 days |  |  |  |  |  |
| Resided in nursing home/long-term care facility prior to admission | 0 (0.0) | 3 (5.2) | 3 (3.4) | 0.55 |  |
| At time of index blood culture collection |  |  |  |  |  |
| Recent surgical procedure | 4 (13.8) | 6 (10.3) | 10 (11.5) | 0.76 |  |
| High-dose steroid use | 4 (13.8) | 6 (10.3) | 10 (11.5) | 0.73 |  |
| Neutropenia (<500 cells/μL) | 11 (37.9) | 22 (37.9) | 33 (37.9) | 1 | |
| Receiving hemodialysis or renal replacement therapy | 6 (20.7) | 6 (10.3) | 12 (13.8) | 0.32 |  |
| Central line placement | 21 (72.4) | 42 (72.4) | 63 (72.4) | 1 |  |
| Urinary catheter | 4 (13.8) | 19 (32.8) | 23 (26.4) | 0.07* | **0.01** |
| Mechanical ventilation | 6 (20.7) | 15 (25.9) | 21 (24.1) | 0.79 |  |
| Pitt bacteremia score $\geq 2$ | 14 (48.3) | 27 (46.6)* | 41 (47.1) | 1 | |
| Index bacteremia episode |  |  |  |  |  |
| Positive culture > 2 days of admission | 16 (55.2) | 29 (50.0) | 45 (51.7) | 0.05* | 0.06 |
| Polymicrobial BSI | 7 (24.1) | 16 (27.6) | 23 (26.1) | 0.93 |  |
| *Enterococcus faecium* | 15 (51.7) | 30 (51.7) | 45 (51.7) | N/A; matching factor |  |
| *Enterococcus faecalis* | 14 (48.3) | 28 (48.3) | 42 (48.3) | N/A; matching factor |  |
| Isolate is vancomycin resistant | 13 (44.8) | 20 (34.5) | 33 (37.9) | 0.48 |  |
| Infectious diseases consult | 28 (96.6) | 49 (84.5) | 77 (88.5) | 0.15 |  |
| Endocarditis | 6 (20.7) | 3 (5.2) | 9 (10.3) | 0.06* | 0.22 |
| Infection source |  |  |  |  |  |
| Central line infection | 5 (17.2) | 8 (13.8) | 13 (14.9) | 0.75 |  |
| Genitourinary | 1 (3.4) | 3 (5.2) | 4 (4.6) | 1 |  |
| Abdominal/gastrointestinal | 9 (31.0) | 13 (22.4) | 22 (25.3) | 0.44 |  |
| Endovascular/endocarditis | 4 (13.8) | 2 (3.4) | 6 (6.9) | 0.09* | 0.52 |
| Unknown/primary source | 8 (27.6) | 28 (48.3) | 36 (41.4) | 0.07* | 0.05 |
| Wound/osteoarticular | 0 (0.0) | 2 (3.4) | 2 (2.3) | 0.60 |  |
| Other | 2 (4.8) | 2 (3.4) | 4 (4.5) | 0.55 |  |
| Clinical outcomes |  |  |  |  |  |
| In-hospital mortality | 12 (41.4) | 12 (20.7) | 24 (27.6) | 0.07* | **0.03** |
| Definitive antimicrobial therapy |  |  |  |  |  |
| *E. faecium* |  |  |  |  |  |
| *Monotherapy* |  |  |  |  |  |
| β-lactams | 8 (27.6) | 8 (13.8) | 16 (18.4) | 0.65 |  |
| Daptomycin | 7 (24.1) | 7 (12.1) | 14 (16.1) | 0.42 |  |
| Vancomycin | 2 (6.9) | 10 (17.2) | 12 (13.8) | 1 |  |
| Linezolid | 2 (6.9) | 5 (8.6) | 7 (8.0) | 1 |  |
| Tigecycline | 3 (10.3) | 1 (1.7) | 4 (4.6) | 0.25 |  |
| *Combination therapy* |  |  |  | 1 |  |
| Dual β-lactams | 3 (10.3) | 7 (12.1) | 10 (11.5) | 1 |  |
| Vancomycin plus β-lactams | 0 (0.0) | 4 (6.9) | 4 (4.6) | 0.54 |  |
| Daptomycin plus β-lactams | 5 (17.2) | 11 (19.0) | 16 (18.4) | 1 |  |
| Daptomycin plus linezolid | 0 (0.0) | 4 (6.9) | 4 (4.6) | 0.54 |  |
| *Other<sup>^</sup>* | 2 (6.9) | 1 (1.7) | 3 (3.4) | 0.25 |  |
| Therapy received was not appropriate | 2 (6.9) | 4 (6.8) | 6 (6.9) | 1 |  |
| *E. faecalis* |  |  |  |  |  |
| *Monotherapy* |  |  |  |  |  |
| β-lactams | 7 (50.0) | 4 (14.3) | 11 (26.2) | **0.04** |  |
| Daptomycin | 1 (7.1) | 0 (0.0) | 1 (2.4) | 0.33 |  |
| Vancomycin | 1 (7.1) | 8 (28.6) | 9 (21.4) | 0.23 |  |
| Linezolid | 0 (0.0) | 1 (3.6) | 1 (2.4) | 1 |  |
| Tigecycline | 1 (7.1) | 0 (0.0) | 1 (2.4) | 0.33 |  |
| *Combination therapy* |  |  |  | 1 |  |
| Dual β-lactams | 3 (21.4) | 7 (25.0) | 10 (23.8) | 1 |  |
| Vancomycin plus β-lactams | 0 (0.0) | 2 (7.1) | 2 (4.8) | 0.54 |  |
| Daptomycin plus β-lactams | 1 (7.1) | 3 (10.7) | 4 (9.5) | 1 |  |
| Daptomycin plus linezolid | 0 (0.0) | 1 (3.6) | 1 (2.4) | 1 |  |
| Therapy received was not appropriate | 0 (0.0) | 6 (21.4) | 6 (14.3) | 0.08* |  |
| Empiric therapy |  |  |  |  |  |
| *E. faecium* |  |  |  |  |  |
| Vancomycin | 2 (13.3) | 5 (16.7) | 7 (15.6) | 1 |  |
| β-lactams | 4 (26.7) | 16 (53.3) | 20 (44.4) | 0.12 |  |
| Daptomycin | 7 (46.7) | 10 (33.3) | 17 (37.8) | 0.59 |  |
| Linezolid | 2 (13.3) | 9 (30.0) | 11 (24.4) | 0.29 |  |
| Tigecycline | 2 (13.3) | 2 (6.7) | 4 (8.9) | 0.59 |  |
| Other<sup>#</sup> | 1 (6.7) | 4 (13.3) | 5 (11.1) | 0.65 |  |
| Therapy received was not appropriate | 5 (33.3) | 2 (6.7) | 7 (15.6) | **0.03** |  |
| *E. faecalis* |  |  |  |  |  |
| Vancomycin | 5 (35.7) | 14 (50.0) | 19 (45.2) | 0.58 |  |
| β-lactams | 4 (28.6) | 13 (46.4) | 17 (40.5) | 0.33 |  |
| Daptomycin | 1 (7.1) | 1 (3.6) | 2 (4.8) | 1 |  |
| Linezolid | 1 (7.1) | 2 (7.1) | 3 (7.1) | 1 |  |
| Tigecycline | 1 (7.1) | 1 (3.6) | 2 (4.8) | 1 |  |
| Other<sup>^^</sup> | 2 (14.3) | 1 (3.6) | 3 (7.1) | 0.25 |  |
| Therapy received was not appropriate | 2 (14.2) | 8 (28.6) | 10 (23.8) | 0.45 |  |

Table values are reported as n(%).
CC- cancer center; HS- hospital system; GLMM- generalized linear mixed model; NA- not applicable
<sup>^</sup>included minocycline (n=4), quinupristin/dalfopristin (n=1); <sup>#</sup>included minocycline (n=5); <sup>^^</sup>included doxycycline (n=1), levofloxacin (n=1), rifampin (n=1)
This table describes clinical characteristics of patients with and without persistent bacteremia.
Matching factors were not analyzed, as they were balanced by design with respect to the outcome of interest. Non-antibiotic variable p-values with an asterisk correspond to variables with p-values above the threshold for significance ( $p < 0.05$ ) but were included in the multivariable analysis. As analyses were performed by species for all antibiotic therapy data, none of these variables were included in the multivariable model.

**Table 2.**
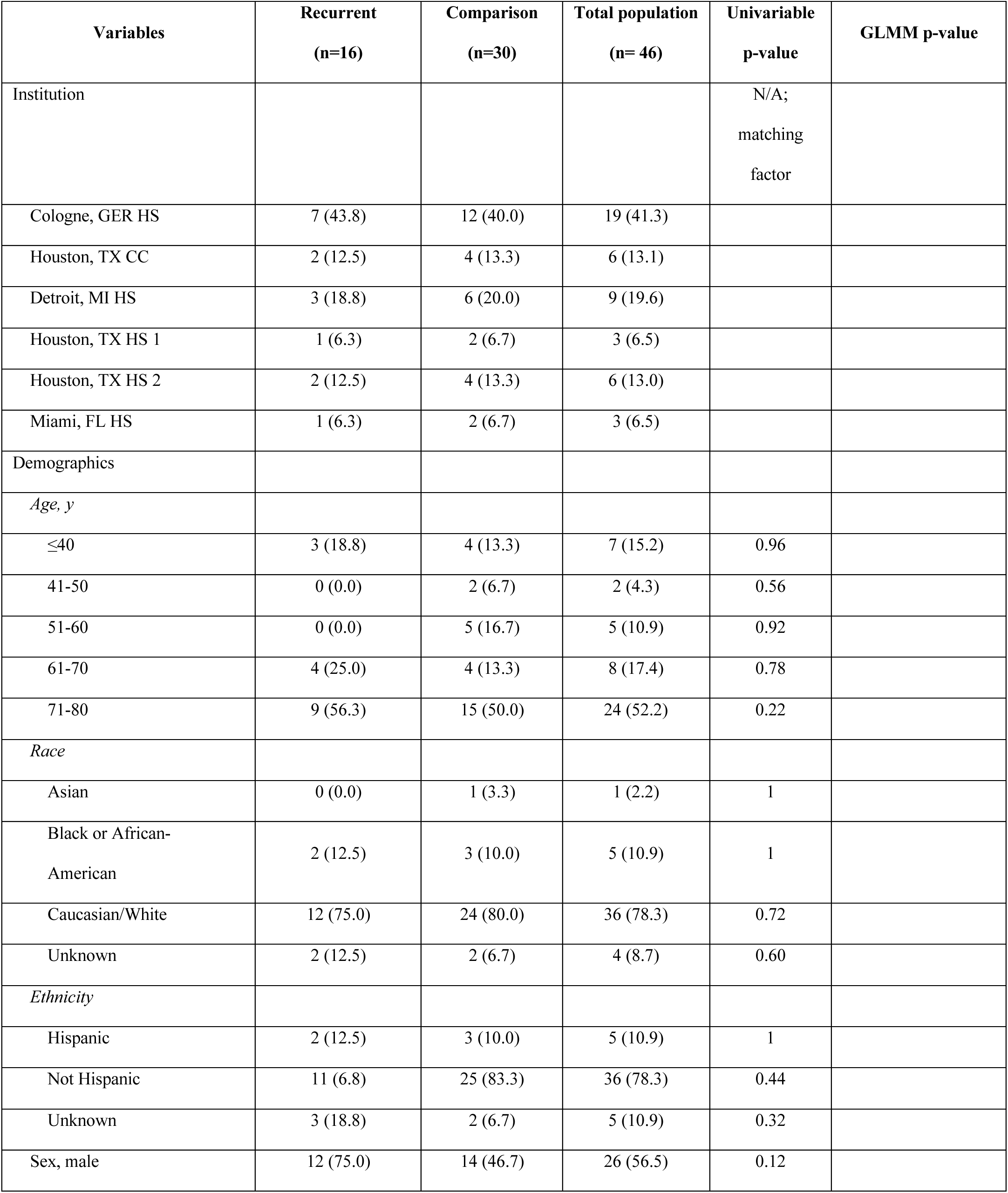
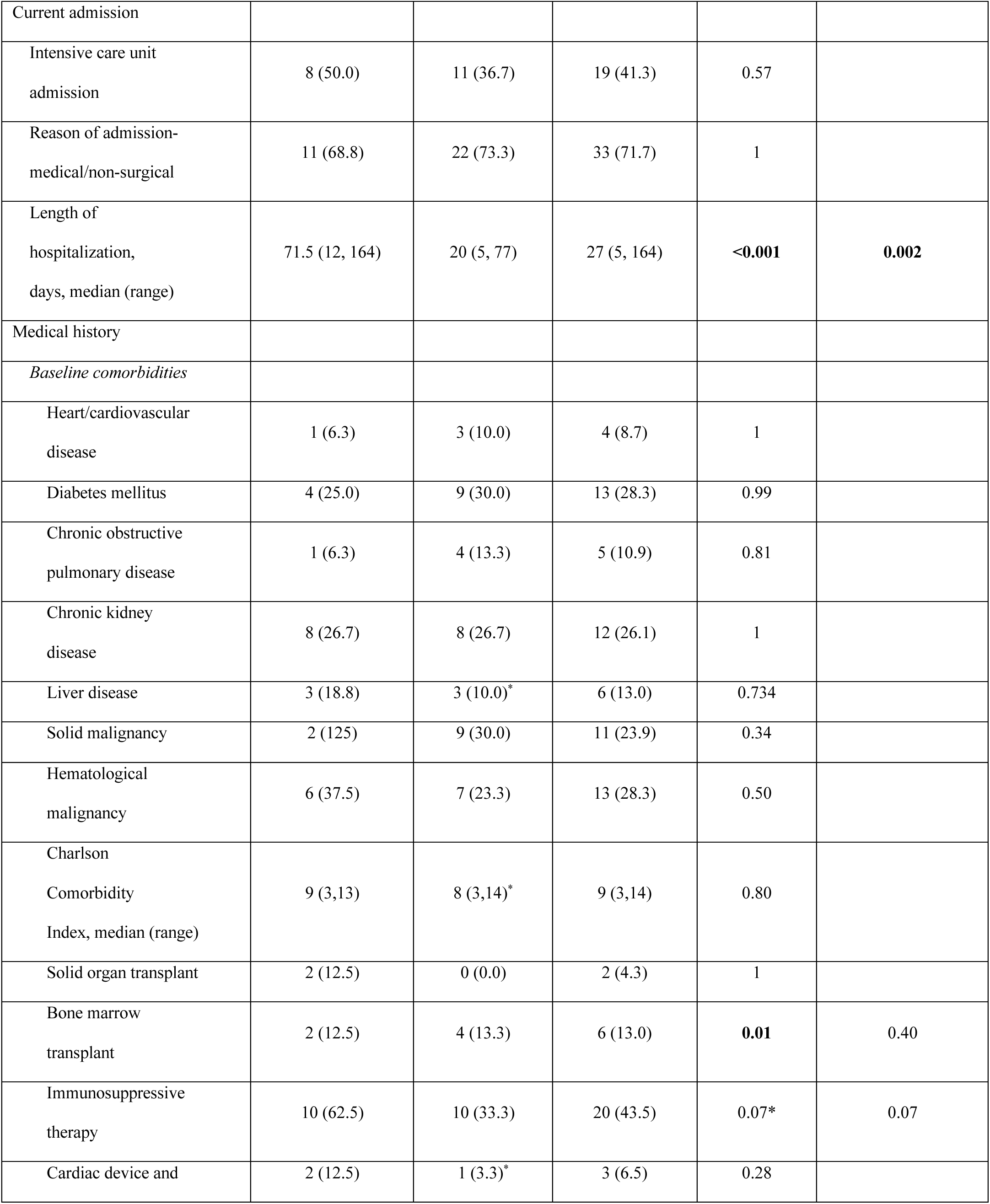
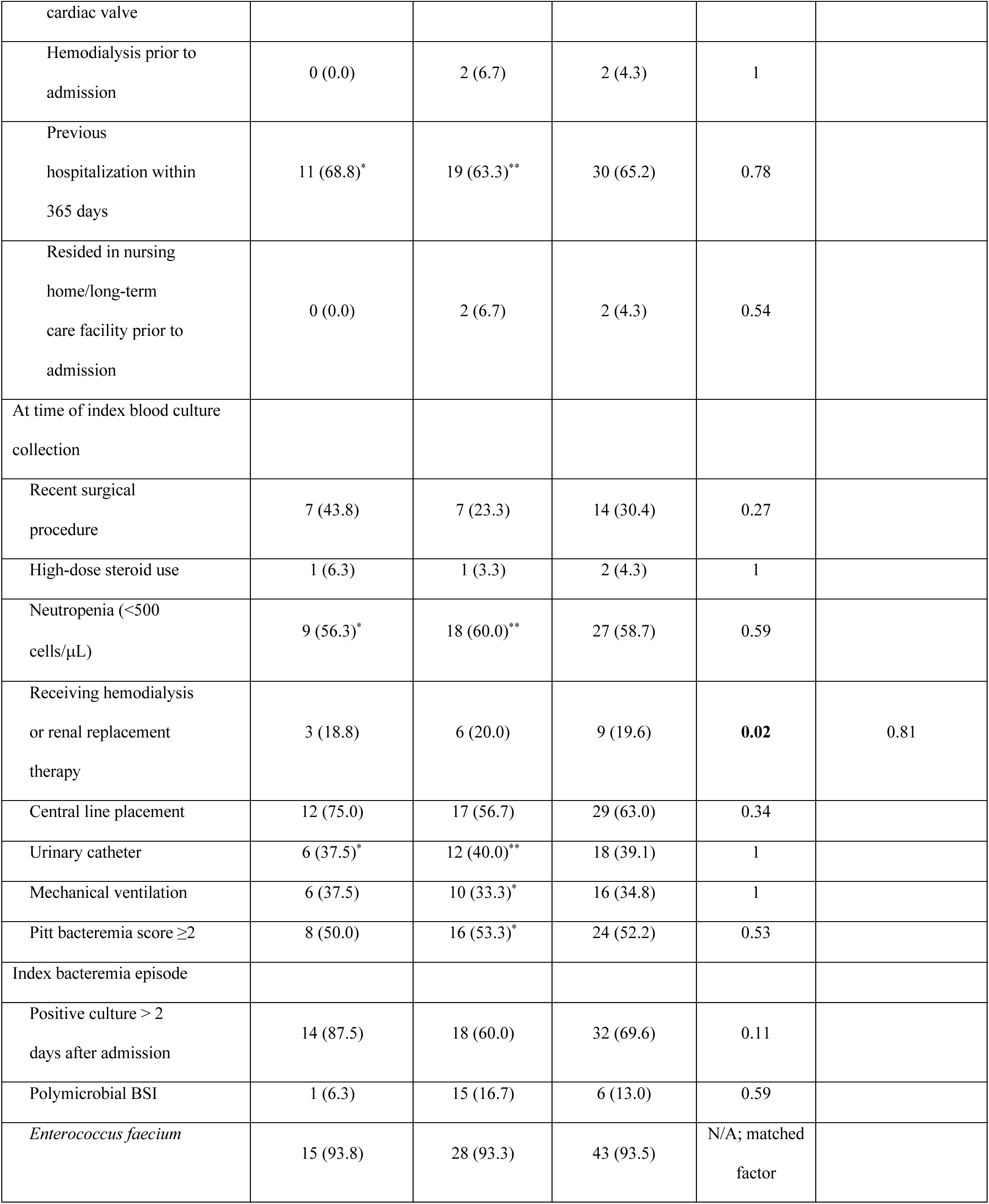
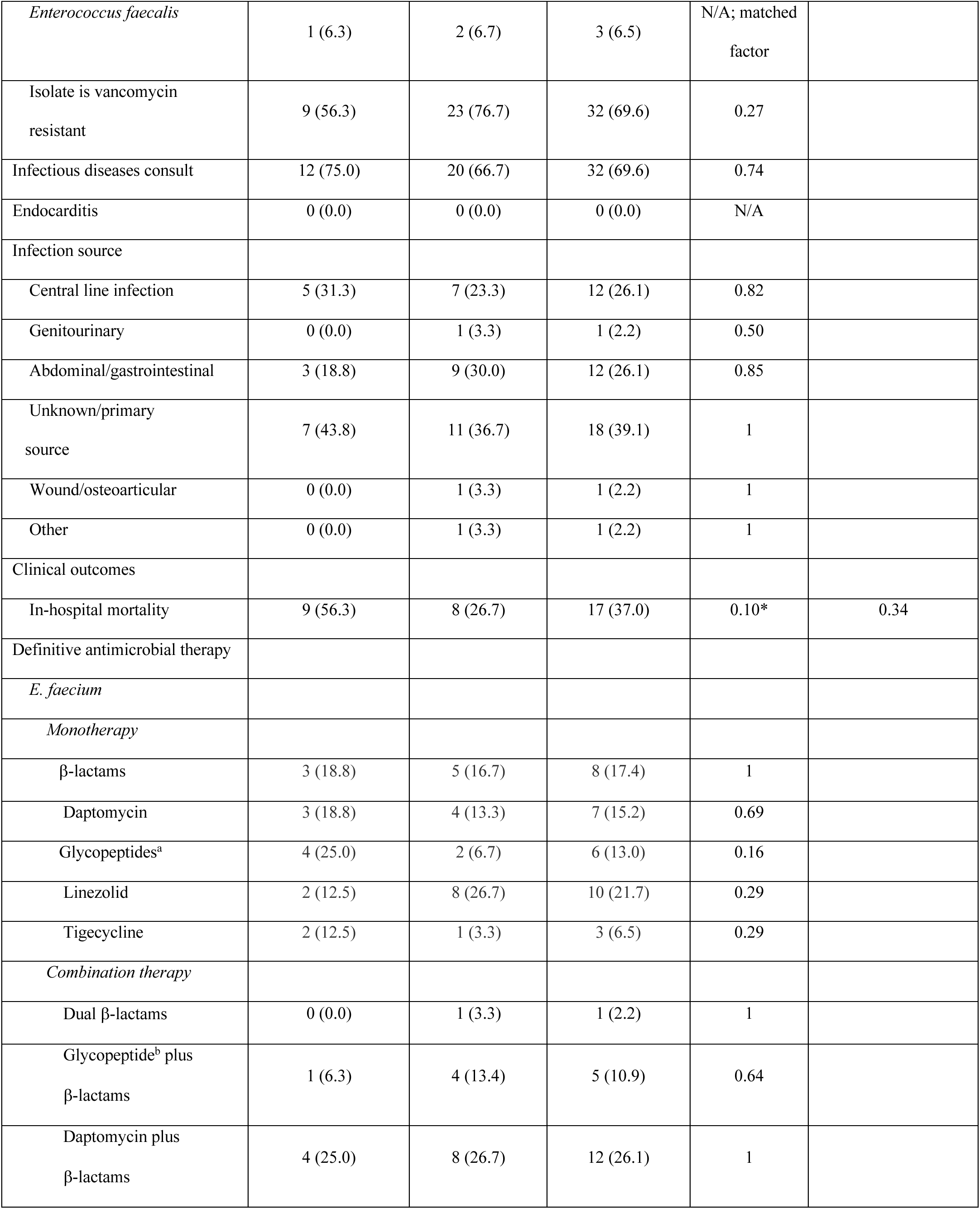
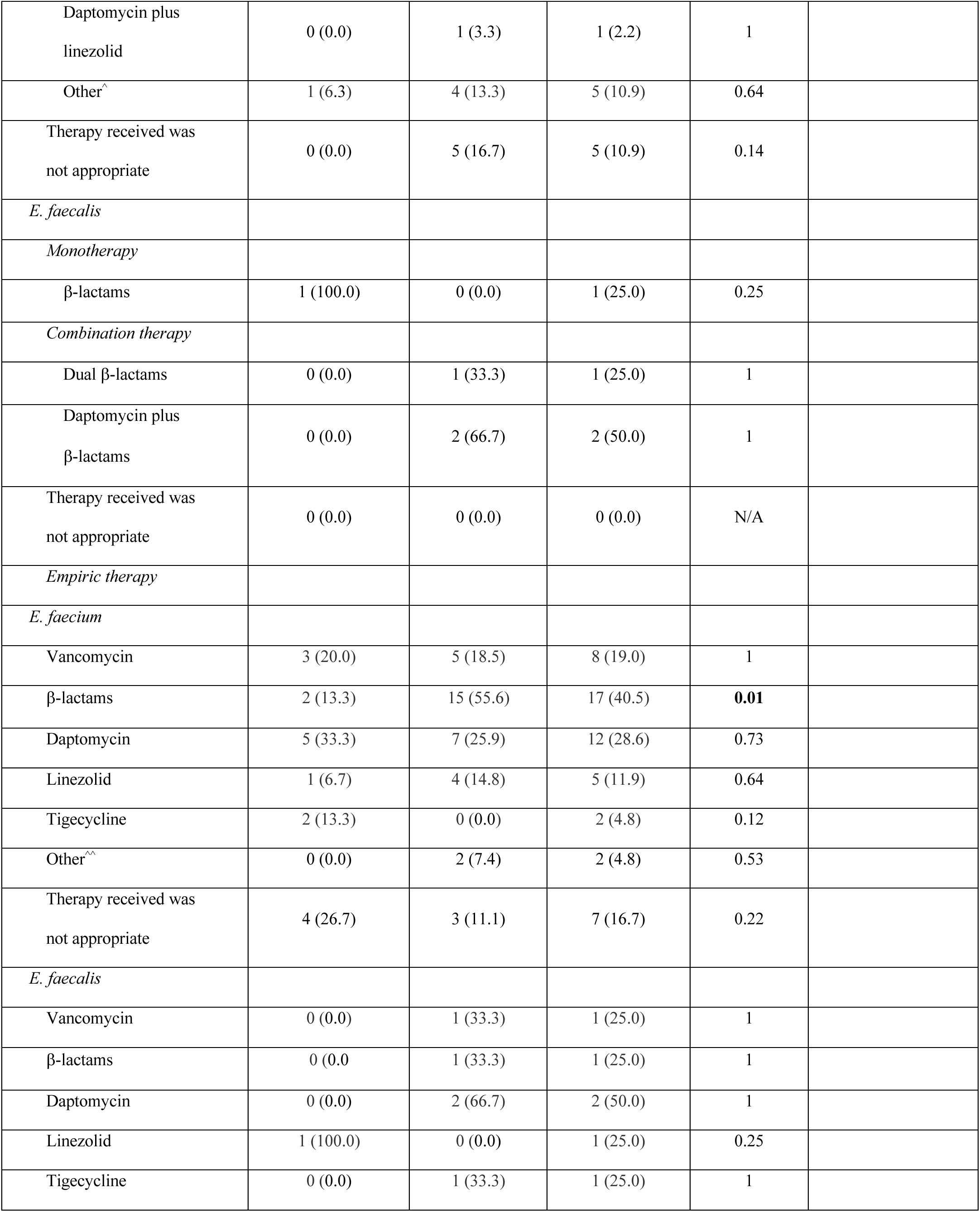
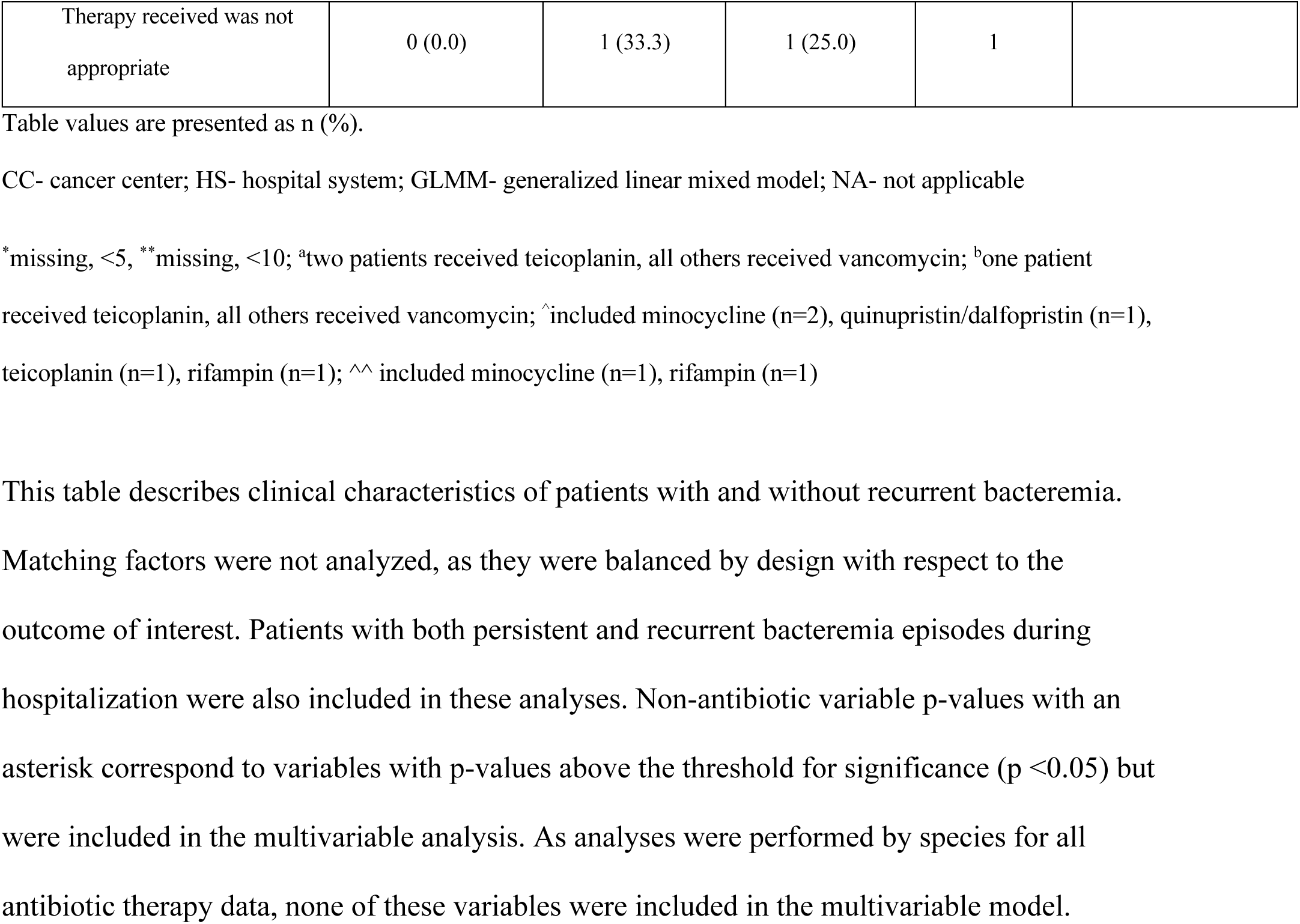
Clinical characteristics of patients with recurrent enterococcal bacteremia and comparison patients.

| Variables | Recurrent<br>(n=16) | Comparison<br>(n=30) | Total population<br>(n= 46) | Univariable<br>p-value | GLMM p-value |
| --- | --- | --- | --- | --- | --- |
| Institution |  |  |  | N/A;<br>matching<br>factor |  |
| Cologne, GER HS | 7 (43.8) | 12 (40.0) | 19 (41.3) |  |  |
| Houston, TX CC | 2 (12.5) | 4 (13.3) | 6 (13.1) |  |  |
| Detroit, MI HS | 3 (18.8) | 6 (20.0) | 9 (19.6) |  |  |
| Houston, TX HS 1 | 1 (6.3) | 2 (6.7) | 3 (6.5) |  |  |
| Houston, TX HS 2 | 2 (12.5) | 4 (13.3) | 6 (13.0) |  |  |
| Miami, FL HS | 1 (6.3) | 2 (6.7) | 3 (6.5) |  |  |
| Demographics |  |  |  |  |  |
| *Age, y* |  |  |  |  |  |
| ≤40 | 3 (18.8) | 4 (13.3) | 7 (15.2) | 0.96 |  |
| 41-50 | 0 (0.0) | 2 (6.7) | 2 (4.3) | 0.56 |  |
| 51-60 | 0 (0.0) | 5 (16.7) | 5 (10.9) | 0.92 |  |
| 61-70 | 4 (25.0) | 4 (13.3) | 8 (17.4) | 0.78 |  |
| 71-80 | 9 (56.3) | 15 (50.0) | 24 (52.2) | 0.22 |  |
| *Race* |  |  |  |  |  |
| Asian | 0 (0.0) | 1 (3.3) | 1 (2.2) | 1 |  |
| Black or African-American | 2 (12.5) | 3 (10.0) | 5 (10.9) | 1 |  |
| Caucasian/White | 12 (75.0) | 24 (80.0) | 36 (78.3) | 0.72 |  |
| Unknown | 2 (12.5) | 2 (6.7) | 4 (8.7) | 0.60 |  |
| *Ethnicity* |  |  |  |  |  |
| Hispanic | 2 (12.5) | 3 (10.0) | 5 (10.9) | 1 |  |
| Not Hispanic | 11 (6.8) | 25 (83.3) | 36 (78.3) | 0.44 |  |
| Unknown | 3 (18.8) | 2 (6.7) | 5 (10.9) | 0.32 |  |
| Sex, male | 12 (75.0) | 14 (46.7) | 26 (56.5) | 0.12 |  |
| Current admission |  |  |  |  |  |
| Intensive care unit admission | 8 (50.0) | 11 (36.7) | 19 (41.3) | 0.57 |  |
| Reason of admission-medical/non-surgical | 11 (68.8) | 22 (73.3) | 33 (71.7) | 1 |  |
| Length of hospitalization, days, median (range) | 71.5 (12, 164) | 20 (5, 77) | 27 (5, 164) | **<0.001** | **0.002** |
| Medical history |  |  |  |  |  |
| *Baseline comorbidities* |  |  |  |  |  |
| Heart/cardiovascular disease | 1 (6.3) | 3 (10.0) | 4 (8.7) | 1 |  |
| Diabetes mellitus | 4 (25.0) | 9 (30.0) | 13 (28.3) | 0.99 |  |
| Chronic obstructive pulmonary disease | 1 (6.3) | 4 (13.3) | 5 (10.9) | 0.81 |  |
| Chronic kidney disease | 8 (26.7) | 8 (26.7) | 12 (26.1) | 1 |  |
| Liver disease | 3 (18.8) | 3 (10.0)* | 6 (13.0) | 0.734 |  |
| Solid malignancy | 2 (12.5) | 9 (30.0) | 11 (23.9) | 0.34 |  |
| Hematological malignancy | 6 (37.5) | 7 (23.3) | 13 (28.3) | 0.50 |  |
| Charlson Comorbidity Index, median (range) | 9 (3,13) | 8 (3,14)* | 9 (3,14) | 0.80 |  |
| Solid organ transplant | 2 (12.5) | 0 (0.0) | 2 (4.3) | 1 |  |
| Bone marrow transplant | 2 (12.5) | 4 (13.3) | 6 (13.0) | **0.01** | 0.40 |
| Immunosuppressive therapy | 10 (62.5) | 10 (33.3) | 20 (43.5) | 0.07* | 0.07 |
| Cardiac device and | 2 (12.5) | 1 (3.3)* | 3 (6.5) | 0.28 |  |
| cardiac valve |  |  |  |  |  |
| Hemodialysis prior to admission | 0 (0.0) | 2 (6.7) | 2 (4.3) | 1 |  |
| Previous hospitalization within 365 days | 11 (68.8)* | 19 (63.3)** | 30 (65.2) | 0.78 |  |
| Resided in nursing home/long-term care facility prior to admission | 0 (0.0) | 2 (6.7) | 2 (4.3) | 0.54 |  |
| At time of index blood culture collection |  |  |  |  |  |
| Recent surgical procedure | 7 (43.8) | 7 (23.3) | 14 (30.4) | 0.27 |  |
| High-dose steroid use | 1 (6.3) | 1 (3.3) | 2 (4.3) | 1 |  |
| Neutropenia (<500 cells/μL) | 9 (56.3)* | 18 (60.0)** | 27 (58.7) | 0.59 | |
| Receiving hemodialysis or renal replacement therapy | 3 (18.8) | 6 (20.0) | 9 (19.6) | **0.02** | 0.81 |
| Central line placement | 12 (75.0) | 17 (56.7) | 29 (63.0) | 0.34 |  |
| Urinary catheter | 6 (37.5)* | 12 (40.0)** | 18 (39.1) | 1 |  |
| Mechanical ventilation | 6 (37.5) | 10 (33.3)* | 16 (34.8) | 1 |  |
| Pitt bacteremia score $\geq 2$ | 8 (50.0) | 16 (53.3)* | 24 (52.2) | 0.53 | |
| Index bacteremia episode |  |  |  |  |  |
| Positive culture > 2 days after admission | 14 (87.5) | 18 (60.0) | 32 (69.6) | 0.11 |  |
| Polymicrobial BSI | 1 (6.3) | 15 (16.7) | 6 (13.0) | 0.59 |  |
| *Enterococcus faecium* | 15 (93.8) | 28 (93.3) | 43 (93.5) | N/A; matched factor |  |
| *Enterococcus faecalis* | 1 (6.3) | 2 (6.7) | 3 (6.5) | N/A; matched factor |  |
| Isolate is vancomycin resistant | 9 (56.3) | 23 (76.7) | 32 (69.6) | 0.27 |  |
| Infectious diseases consult | 12 (75.0) | 20 (66.7) | 32 (69.6) | 0.74 |  |
| Endocarditis | 0 (0.0) | 0 (0.0) | 0 (0.0) | N/A |  |
| Infection source |  |  |  |  |  |
| Central line infection | 5 (31.3) | 7 (23.3) | 12 (26.1) | 0.82 |  |
| Genitourinary | 0 (0.0) | 1 (3.3) | 1 (2.2) | 0.50 |  |
| Abdominal/gastrointestinal | 3 (18.8) | 9 (30.0) | 12 (26.1) | 0.85 |  |
| Unknown/primary source | 7 (43.8) | 11 (36.7) | 18 (39.1) | 1 |  |
| Wound/osteoarticular | 0 (0.0) | 1 (3.3) | 1 (2.2) | 1 |  |
| Other | 0 (0.0) | 1 (3.3) | 1 (2.2) | 1 |  |
| Clinical outcomes |  |  |  |  |  |
| In-hospital mortality | 9 (56.3) | 8 (26.7) | 17 (37.0) | 0.10* | 0.34 |
| Definitive antimicrobial therapy |  |  |  |  |  |
| *E. faecium* |  |  |  |  |  |
| *Monotherapy* |  |  |  |  |  |
| β-lactams | 3 (18.8) | 5 (16.7) | 8 (17.4) | 1 |  |
| Daptomycin | 3 (18.8) | 4 (13.3) | 7 (15.2) | 0.69 |  |
| Glycopeptides<sup>a</sup> | 4 (25.0) | 2 (6.7) | 6 (13.0) | 0.16 |  |
| Linezolid | 2 (12.5) | 8 (26.7) | 10 (21.7) | 0.29 |  |
| Tigecycline | 2 (12.5) | 1 (3.3) | 3 (6.5) | 0.29 |  |
| *Combination therapy* |  |  |  |  |  |
| Dual β-lactams | 0 (0.0) | 1 (3.3) | 1 (2.2) | 1 |  |
| Glycopeptide<sup>b</sup> plus β-lactams | 1 (6.3) | 4 (13.4) | 5 (10.9) | 0.64 |  |
| Daptomycin plus β-lactams | 4 (25.0) | 8 (26.7) | 12 (26.1) | 1 |  |
| Daptomycin plus<br>linezolid | 0 (0.0) | 1 (3.3) | 1 (2.2) | 1 |  |
| Other<sup>^</sup> | 1 (6.3) | 4 (13.3) | 5 (10.9) | 0.64 |  |
| Therapy received was<br>not appropriate | 0 (0.0) | 5 (16.7) | 5 (10.9) | 0.14 |  |
| *E. faecalis* |  |  |  |  |  |
| *Monotherapy* |  |  |  |  |  |
| β-lactams | 1 (100.0) | 0 (0.0) | 1 (25.0) | 0.25 |  |
| *Combination therapy* |  |  |  |  |  |
| Dual β-lactams | 0 (0.0) | 1 (33.3) | 1 (25.0) | 1 |  |
| Daptomycin plus<br>β-lactams | 0 (0.0) | 2 (66.7) | 2 (50.0) | 1 |  |
| Therapy received was<br>not appropriate | 0 (0.0) | 0 (0.0) | 0 (0.0) | N/A |  |
| *Empiric therapy* |  |  |  |  |  |
| *E. faecium* |  |  |  |  |  |
| Vancomycin | 3 (20.0) | 5 (18.5) | 8 (19.0) | 1 |  |
| β-lactams | 2 (13.3) | 15 (55.6) | 17 (40.5) | **0.01** |  |
| Daptomycin | 5 (33.3) | 7 (25.9) | 12 (28.6) | 0.73 |  |
| Linezolid | 1 (6.7) | 4 (14.8) | 5 (11.9) | 0.64 |  |
| Tigecycline | 2 (13.3) | 0 (0.0) | 2 (4.8) | 0.12 |  |
| Other<sup>^^</sup> | 0 (0.0) | 2 (7.4) | 2 (4.8) | 0.53 |  |
| Therapy received was<br>not appropriate | 4 (26.7) | 3 (11.1) | 7 (16.7) | 0.22 |  |
| *E. faecalis* |  |  |  |  |  |
| Vancomycin | 0 (0.0) | 1 (33.3) | 1 (25.0) | 1 |  |
| β-lactams | 0 (0.0) | 1 (33.3) | 1 (25.0) | 1 |  |
| Daptomycin | 0 (0.0) | 2 (66.7) | 2 (50.0) | 1 |  |
| Linezolid | 1 (100.0) | 0 (0.0) | 1 (25.0) | 0.25 |  |
| Tigecycline | 0 (0.0) | 1 (33.3) | 1 (25.0) | 1 |  |
| Therapy received was not appropriate | 0 (0.0) | 1 (33.3) | 1 (25.0) | 1 |  |

Table values are presented as n (%).
CC- cancer center; HS- hospital system; GLMM- generalized linear mixed model; NA- not applicable
\*missing, <5, \*\*missing, <10; <sup>a</sup>two patients received teicoplanin, all others received vancomycin; <sup>b</sup>one patient received teicoplanin, all others received vancomycin; <sup>^</sup>included minocycline (n=2), quinupristin/dalfopristin (n=1), teicoplanin (n=1), rifampin (n=1); <sup>^^</sup> included minocycline (n=1), rifampin (n=1)
This table describes clinical characteristics of patients with and without recurrent bacteremia.
Matching factors were not analyzed, as they were balanced by design with respect to the outcome of interest. Patients with both persistent and recurrent bacteremia episodes during hospitalization were also included in these analyses. Non-antibiotic variable p-values with an asterisk correspond to variables with p-values above the threshold for significance ( $p < 0.05$ ) but were included in the multivariable analysis. As analyses were performed by species for all antibiotic therapy data, none of these variables were included in the multivariable model.

Patients with persistent vs. non-persistent bacteremia differed in that they were predominately male, more often admitted to the ICU upon admission, and had longer hospital stays following collection of the index positive blood culture. There was no difference between persistent and comparison groups regarding relevant comorbidities, including neutropenia, solid organ transplant history, or presence of hematological malignancies (**Table 1**), though a larger proportion of patients with persistent bacteremia had endocarditis during hospitalization relative to the comparison group. In the multivariable model, length of hospitalization, presence of a urinary catheter at the time of index blood culture, and in-hospital mortality were significantly associated with persistent bacteremia. Notably, a higher proportion of patients with persistent *E. faecium* bacteremia received inappropriate empiric therapy relative to those in the comparison group (5/15, 33.3% vs. 2/30, 6.7%, respectively) (**Table 1**).

Patients with recurrent bacteremia had increased lengths of hospitalization following the index positive blood culture (median, recurrent: 71.5 days; vs. 20 days, p < 0.001), were more likely to have a history of bone marrow transplant, and be actively receiving renal replacement therapy relative to the comparison group (**Table 2**). However, only length of hospitalization was significantly associated with recurrent bacteremia in the adjusted analysis (**Table 2**). A composite analysis of all recalcitrant (persistent and recurrent) bacteremia patients indicated that those with recalcitrant bacteremia were more likely to be male, have a longer hospital stay following index culture, receive immunosuppressive therapy at baseline, and have endocarditis during hospitalization compared to those without recalcitrant bacteremia (**Supplementary Material, Table S1**).

Overall, there was no predominating sequence type (ST) associated with persistent or recurrent infections in either *E. faecium* or *E. faecalis* (**Figure 3**). However, there was evidence of institution-specific clustering by sequence types, such as *E. faecium* ST80 and 117 in Cologne, Germany, and *E. faecalis* ST6 in a cancer center in Houston, TX. Nine sequence types were represented in the persistent *E. faecium* cohort (**Figure 3A**) with four unique STs (1471, 1478, 584, and 736). For *E. faecalis*, ST19 and 480 were unique to the persistent cohort (**Figure 3B**). All but two patients in the persistent cohort were consistently infected with the same enterococcal strain, and they were all of the same sequence type (**Supplementary Material, Figure S1**). In the recurrent cohort, the infecting isolates also belonged to the same strain in most cases, with the exception of two patients from Cologne, Germany (ST117 to ST80; >7000 SNP differences) (**Supplementary Material**, **Figures S1, S2**) with changes in vancomycin susceptibility. Extensive recombination with a resulting pairwise SNP distance of >1,000 was found in one patient (**Supplementary Material, Figure S3**).

**Figure 3.**
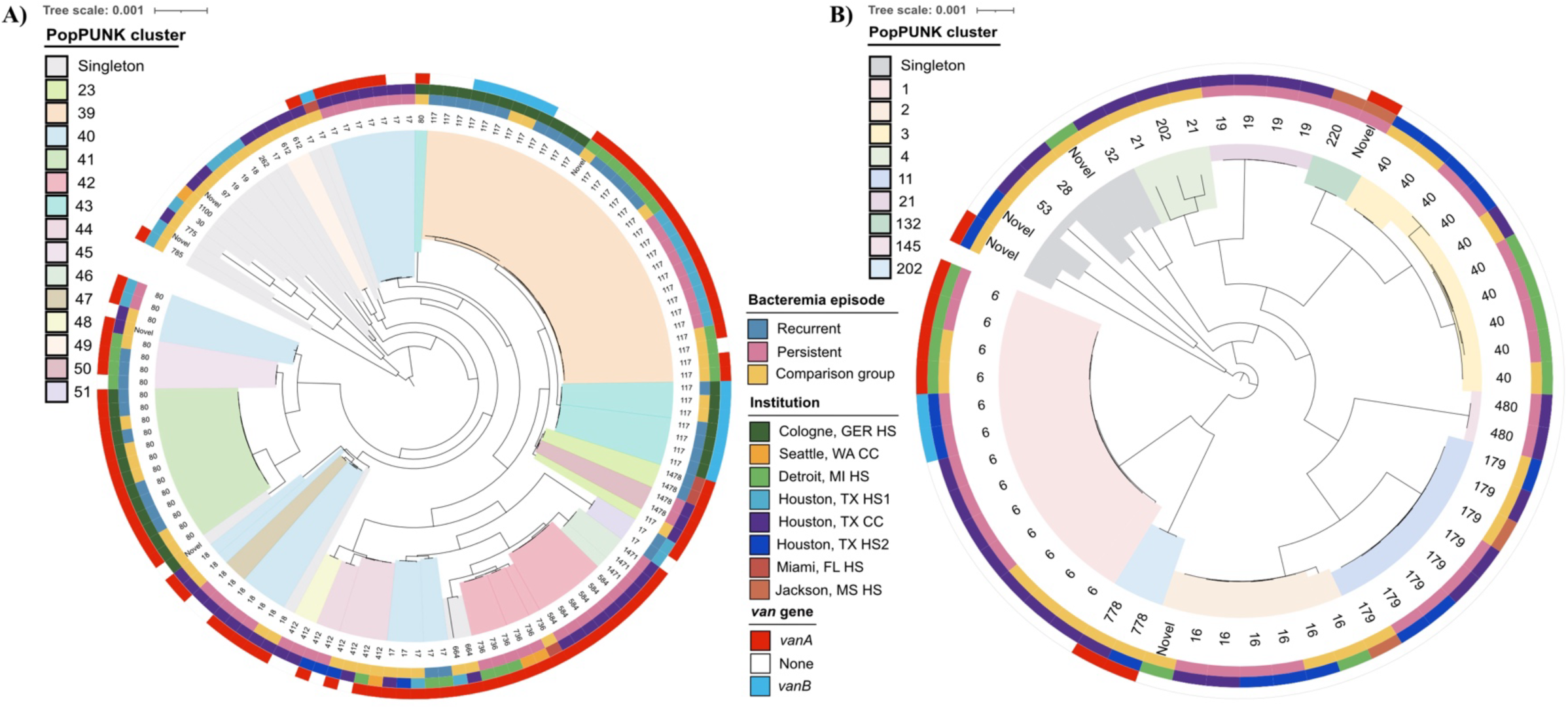
Population structure of the recalcitrant enterococcal bacteremia cohort. (A) *E. faecium* (n=131). (B) *E. faecalis* (n=57). These figures depict midpoint-rooted, core genome-based maximum likelihood phylogenetic trees for comparison, persistent, and recurrent VENOUS isolates. The labels indicate the sequence type (ST) of each isolate, while branch shading indicates PopPUNK cluster, a whole-genome clustering method, as a higher-resolution clustering method relative to ST assignment. Presence or absence of vancomycin resistance genes *vanA* and *vanB*, institution of isolate collection, and bacteremia episode designation are represented as colored bars on the outside of each tree.

To assess the potential of pre-adaptation in isolates causing infection, isolates from the index positive blood culture in the persistent and comparison groups were compared to assess genomic differences (**Supplementary Material**, **Table S2**). A total of 11/15 (73.3%) persistent *E. faecium* index isolates harbored *vanA* relative to 16/30 (46.7%) non-persistent index isolates. Further, persistent *E. faecium* isolates were resistant to more antibiotic classes than their respective comparison cohort, although the number of antimicrobial resistance (AMR) determinants was not significantly different between the groups. Persistent *E. faecium* index isolates also had larger genomes relative to non-persistent index isolates (+85,181 base pairs), including accessory genomes (37.8% accessory gene content vs. 35.7%, respectively), although the median number of plasmids per isolate was identical. However, one small plasmid, *rep* type rep 14b_2, was found in two persistent *E. faecium* index isolates but was absent from non-persistent index isolates (**Supplementary Material, Table S3A**).

In *E. faecalis*, the differences were more modest than those observed in *E. faecium*, but there was a larger proportion of *vanA*-harboring non-persistent index isolates compared to the persistent group [5/28 (17.9%) vs. 1/14 (7.1%)]. Of note, one index ST6 *E faecalis* isolate from a Houston patient with persistent bacteremia harbored *vanB*, a previously published occurrence.^13^ Persistent *E. faecalis* isolates did not harbor significantly different numbers of AMR determinants, nor were they resistant to more antibiotic classes relative to controls(**Supplementary Material, Table S2B**). Lastly, one plasmid *rep* type, rep11b_1, was present in two *E. faecalis* index isolates from the persistent bacteremia group but absent in non-persistent index isolates (**Supplementary Material, Table S3B; Figure S4**). This 9,331-bp plasmid has been characterized and contains enterocins 1071A and 1071B.^14^

Next, the average proportions of functional gene classes (Clusters of Orthologous Groups, or COGs) present in the accessory genomes of persistent vs. non-persistent index isolates were compared for *E. faecium* and *E. faecalis* (**Supplementary Material, Figure S5**). There were no significant differences in proportions of COG groups between persistent and non-persistent *E. faecium* index isolates, although a larger proportion of genes belonging to COG L (replication, recombination, and repair) was found in persistent index isolates (**Figure 4A**). Further examination of gene groups exclusively found in persistent isolates (n=703) revealed 18 gene groups that were significantly associated with persistence, half of which belonged to COG L and were predominantly transposases (**Supplementary Data 2,** tabs 2, 3). In *E. faecalis*, there was a significantly larger proportion of genes belonging to COG G, associated with carbohydrate transport and metabolism (**Figure 4B**). However, of gene groups exclusively present in persistent isolates (n= 646), only one, a gene associated with a phage protein (group_5172), was significantly associated with persistent BSI and could not be assigned to any COG functional group (**Supplementary Data 2,** tabs 3, 4).

**Figure 4.**
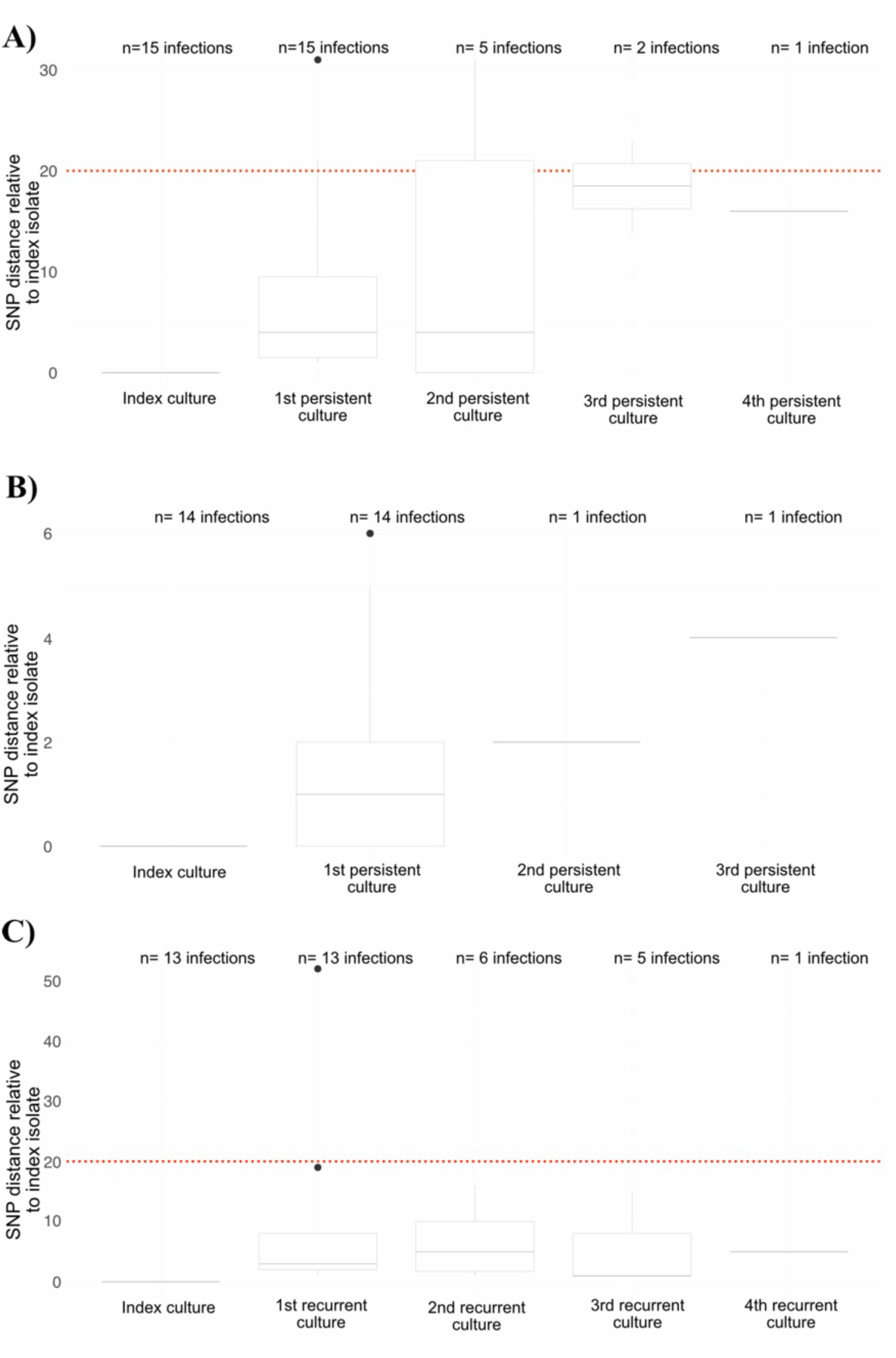
SNP distances relative to index culture of recalcitrant isolates by time of culture collection. (A) Box plot of pairwise SNP distance relative to index culture isolate for persistent *E.* faecium infections; (B) Box plot of pairwise SNP distances relative to the index culture isolate for persistent *E. faecalis* infections; (C) Box plot of pairwise SNP distances relative to the index culture isolate for recurrent *E. faecium* infections. The red dotted line corresponds to the previously published 20-SNP threshold for strain differentiation. For recurrent infections, three infections were excluded from these analyses—two due to ST switching, and one due to a SNP distance of >1000 due to disruptive genomic insertions.

Subsequently, we assessed genetic changes during each persistent infection from the index culture to the sequential persistent positive culture to evaluate whether positive genetic selection occurred in response to host or antibiotic pressures. Median genome size of *E. faecium* isolates decreased over time by 50,808 base pairs, although the median number of plasmids stayed the same (n=4). Gains or losses in the typeable plasmid repertoire were observed in five persistent bacteremia episodes (5/15; 33.3%) (**Supplementary Material**, **Figure S6A**). In persistent *E. faecalis* infections, genomic changes were smaller relative to *E. faecium*. There was a slight increase (2,261 base pairs) in average genome size over time, and median plasmid carriage remained the same, though it was overall lower than in *E. faecium* isolates (median carriage of 1 vs. 4, respectively) (**Supplementary Material; Table S2**). However, as observed in *E. faecium*, there were also five plasmid gain/loss events (5/14; 35.7%) observed in the typeable plasmid repertoire in this cohort (**Supplementary Material**, **Figure S6B**). For both species, plasmid gains and losses were not specific to a specific *rep* type.

Chromosomal structural variations (SV) have been postulated to drive niche adaptation through transcriptional regulation and subsequent alteration of bacterial phenotype.^15^ Chromosomal SV were highly prevalent, with SV>1,000 bp occurring in 21/29 (72.4%) infection pairs, and they were observed more often in *E. faecium*. The most common SV observed in *E. faecium* infection pairs was the gain or loss of insertion sequences in ISL3, IS3, or IS256 families. In *E. faecalis* infection pairs, SV predominantly occurred in site-specific recombination regions (**Supplementary Data 2**, tab 11), likely modulating transcription of downstream loci.^15^ A notable SV in *E. faecium* included an IS1216E-mediated insertion of the *vanA* operon and a chromosomal insertion of a cadmium resistance gene *cadC* disrupting *fhs* (formyltetrahydrofolate synthetase gene) (**Supplementary Material, Figure S7**). In *E. faecalis,* a five-fold duplication of a gene cluster that includes a putative multidrug resistance protein (EF-1078) and streptomycin adenylyltransferase gene (*ant1*) was seen in the index isolate, which was reduced to a single copy in the persistent isolate (**Supplementary Material, Figure S8**).

Given the relatively short time between blood cultures, there was no appreciable difference in the proportions of accessory genes belonging to any COG functional categories for either species in each patient’s index blood culture isolate and the respective persistent isolate, suggesting that the persistence of bacteremia is not generally driven by the acquisition of specific genes. However, the proportion of genes belonging to COG L (replication, recombination, repair) was higher in persistent *E. faecium* isolates relative to persistent *E. faecalis* (**Supplementary Material, Figure S9**). Relative to the index blood culture isolate, persistent *E. faecium* isolates gained substantially more genes than *E. faecalis* isolates (median genes gained: 34 vs. 10.5, respectively) **(Supplementary Data 2**, tabs 7, 8). For both species, the majority of genes gained in more than one patient isolate set were associated with COG L group, including transposases from the IS66 family, integrase core domains, and phage integrases. **(Supplementary Data 2,** tabs 9, 10).

Index isolates from persistent bacteremia episodes served as patient-specific references to examine within-host diversity in persistent timepoint isolates relative to baseline (**Figure 4**). Overall, the average pairwise SNP distance was markedly higher for *E. faecium* relative to *E. faecalis* [average: 7.73 (range: 0-31) vs 1.71 (0-6), respectively], with only two infection pairs from both *E. faecium* and *E. faecalis* infections differing by over 20 SNPs (31 and 21 SNPs), the previously published threshold suggested to differentiate *E. faecium* strains ^16^. Nonetheless, 3/5 *E. faecium* persistent infections with more than one positive culture beyond the first persistent time point exceeded this threshold at some point during infection (**Figure 4A**). The overall number of non-synonymous SNPs observed in *E. faecium* was higher compared to *E. faecalis* persistent infections. However, the proportion of non-synonymous SNPs relative to synonymous SNPs was much higher in *E. faecalis* infection pairs (**Figure 5; Supplementary Material,** p 23).

**Figure 5.**
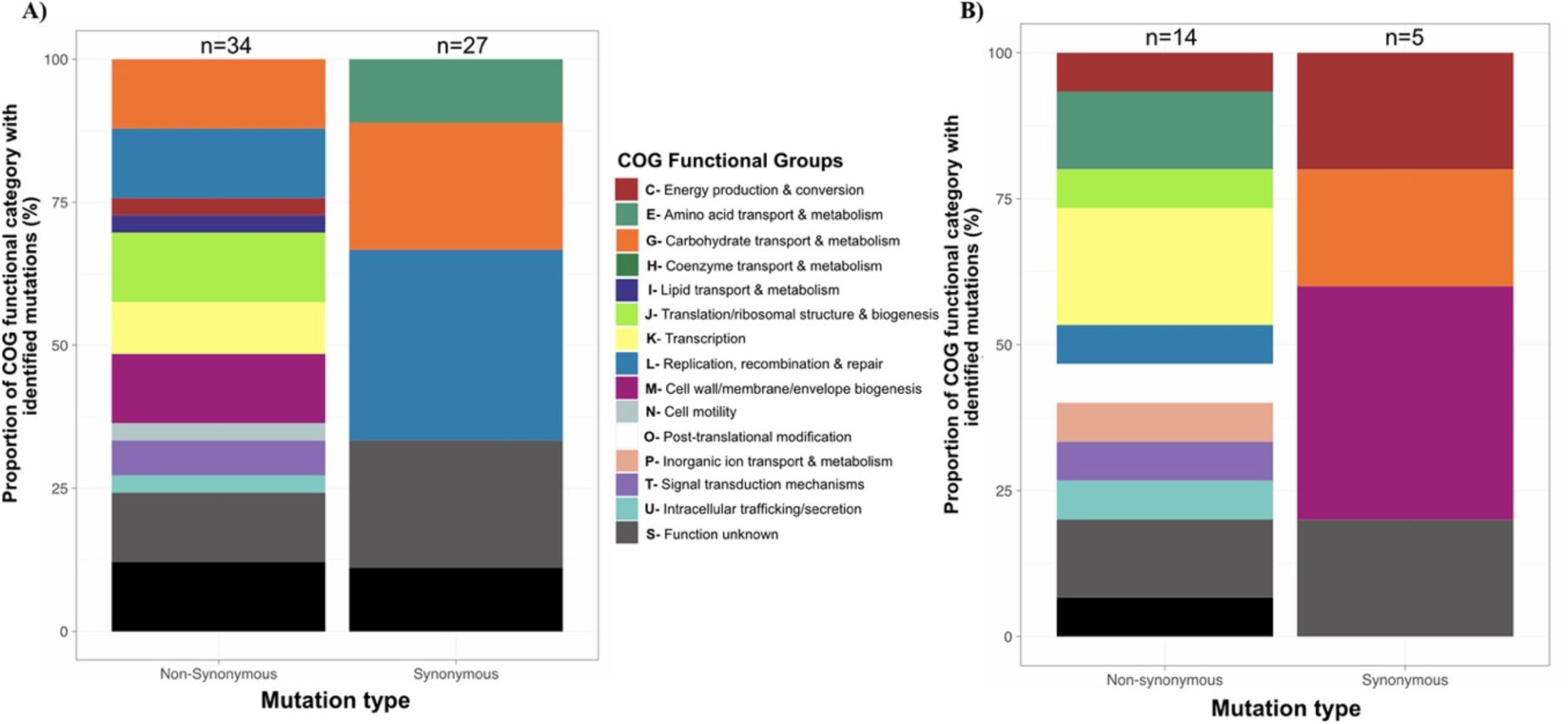
Functional characterization of mutations in coding regions during the course of persistent bacteremia. (A) *E. faecium.* (B) *E. faecalis*. These stacked bar plots indicate the average proportions of the functional classifications of mutation-containing genes in index bacteremia isolates vs. respective persistent time point bacteremia isolates. Synonymous and non-synonymous mutations are presented separately for each species, and the total number of mutations identified in coding regions are presented at the top of each stacked bar plot.

The representation of functional groups associated with mutated genes was relatively disparate between *E. faecium* and *E. faecalis* infection pairs, suggesting species-specific pathways of adaption. For instance, non-synonymous mutations in genes belonging to COG O (post-translational modification) were exclusively found in *E. faecalis* infection pairs (**Figure 5B**). In contrast, there was a markedly higher proportion of mutations in genes belonging to COGs L, (carbohydrate transport/utilization), G (amino acid transport/utilization) and J (translation/ribosomal structure/biogenesis) in *E. faecium* pairs (**Figure 5A**). Of note, a non-synonymous mutation in *rpoB* (resulting in H486N), observed in one *E. faecium* persistent pair, has been associated with rifampin and daptomycin resistance.^17^ Moreover, mutation-based adaptation appears to be episode-specific in this cohort, as there were no common mutations across more than one infection episode in either species. Nonetheless, the same variant, Thr167Ile in a GTP diphosphokinase (encoded by *relA*) and previously associated with the ppGpp stringent response under stress conditions,^18^ appeared in one infection pair in both *E. faecium* and *E. faecalis* (**Supplementary Data 2,** tabs 12, 14).

Two episodes of *E. faecium* recurrent bacteremia exhibited strain switching, exhibiting SNP distances of >20 as well as a change in sequence type. In the other 11 episodes, pairwise SNP distances in recurrent bacteremia infection pairs were significantly higher compared to pairwise SNP distances in persistent *E. faecium* pairs (**Supplementary Material, Figures S10, S11**). Recurrent *E. faecium* isolates also exhibited a larger proportion of non-synonymous mutations relative to the persistent *E. faecium* cohort (**Supplementary Material, Figure S10**). As previously described, one same-ST recurrent bacteremia episode was not included in aggregate analyses— though no strain switching was observed, the recurrent isolate collected differed from the index isolate by 1,462 SNPs, indicating a different infecting strain. Of the SNPs in coding regions, 88.8% (1,030) were located in the chromosome. Further genomic characterization revealed the insertion of large genomic elements in the recurrent isolate (**Supplementary Material, Figures S2, S3**), including a 43.3kb transposon harboring the *vanB* operon that resulted in conversion to phenotypic vancomycin resistance.

## DISCUSSION

In this large-scale, multicenter cohort study of 1,265 patients with enterococcal bacteremia, we found that 41/1,265 (3.2%) of patients experienced a recalcitrant episode. Recalcitrant episodes were further divided into patients with persistent bacteremia (defined as positive blood cultures ≥ 4 days after the index culture) and those with recurrent bloodstream infection (patients who had initial clearance but their bacteremia returned). Our findings indicate that patients with extended hospitalizations or presence of a urinary catheter (likely reflecting chronic illness) are at higher risk of developing persistent bacteremia. Moreover, those patients experienced increased risk of mortality, confirming previous findings using the VENOUS cohort^2^ and supporting the notion that efforts to clear the bacteremia are of paramount importance to improve patients’ outcomes. Of note, there was a relatively even number of persistent *E. faecium* and *E. faecalis* infections, despite the fact that persistent *E. faecalis* were predominantly ampicillin and vancomycin-susceptible and harbored a lower number of AMR determinants than *E. faecium* infections (2 vs. 11.5, respectively), and were most likely to receive appropriate therapy. However, species-specific comparisons showed that nearly 30% of patients with persistent *E. faecalis* infections were also diagnosed with endocarditis during hospitalization, relative to only 13.3% of patients with persistent *E. faecium* infections. Previous studies assessing risk factors for persistent enterococcal bacteremia have found that infective endocarditis is an independent risk factor for persistent bacteremia despite effective therapy.^19^

The only clinical variable significantly associated with recurrent bacteremia was length of hospitalization. The recurrent episodes were caused predominantly by *E. faecium* and were primarily attributed to a single site. This is likely due to the greater frequency of blood cultures per patient taken by this site—nearly daily in most instances— which may have allowed for the identification of clearance events that would not be captured by less frequent blood cultures. Nonetheless, our findings suggest that both phenomena (persistence and recurrence) are clinically distinct and likely are influenced by different factors.

Our comprehensive genomic analyses suggest that persistent *E. faecium* enterococcal bacteremia isolates may be pre-adapted to cause recalcitrant infection and that pathways of adaptation are both species- and episode-specific. More importantly, we found that most episodes of persistent and recurrent infection were with the same enterococcal strain. Our findings suggest that isolates likely evolve independently of lineage to cause persistent infection. Indeed, vancomycin resistance was more prevalent in the persistent group, genome size was markedly larger, and there was resistance to significantly more antibiotic classes relative to non-persistent isolates.

Persistent *E. faecium* isolates had a higher proportion of genes associated with replication, recombination, and repair, and half of the gene groups found exclusively in persistent *E. faecium* isolates were transposases and insertion sequences, consistent with previous findings.^20^ Of note, persistent *E. faecium* isolates exhibited a median decrease in genome size of over 50,000 base pairs despite maintaining a stable median plasmid carriage, indicating that a major source of genetic variation is chromosomal SV. Larger SV of >1000 base pairs were highly prevalent in both *E. faecium* and *E. faecalis* infection pairs, but the type of SVs observed appeared to be largely species-specific, with *E. faecium* SV driven mainly by insertion sequence gain and loss, whereas *E. faecalis* SV were more often attributed to reshuffling or inversion of existing genetic material at site-specific recombination regions. This phenomenon of “phase variation” in *E. faecalis* affecting Type I restriction-modification systems impacts gene expression,^15,21^ while the lack of functional CRISPR-cas systems and highly plastic genomes of clinical *E. faecium* isolates are widely associated with acquisition and loss of mobile genetic elements, such as insertion sequences.^22^

Although *E. faecium* gained over three times more genes during infection relative to *E. faecalis,* ^23^ we found no shared patterns of adaptation, with the exception of a *relA* mutation in one *E. faecium* and one *E. faecalis* infection pair. Mutations in *relA*, a mediator of the bacterial stringent response, have been associated with antibiotic resistance in multiple pathogens, with mutations likely to alter the expression of genes related to biofilm-dependent antibiotic tolerance in immunocompromised hosts.^24^

Despite the large sample of sites and patients compared with previous studies, statistical power is limited due to the relatively small number of recalcitrant episodes. There is also the potential for selection bias, as the distribution of vancomycin resistance in *E. faecalis* isolates in this study does not reflect the overall prevalence of vancomycin-resistant *E. faecalis* in the general U.S. clinical population. Although U.S. geographical representation is robust, the global generalizability of this work is limited given the single non-U.S. study site, especially considering that the infections from this site were exclusively classified as recurrent bloodstream infections. Lastly, only one isolate per blood culture was sequenced, meaning the full breadth of within-host diversity over time may have been missed. However, previous studies have shown that even sequencing up to ten isolates per culture does not reveal substantial diversity in infections from this source.^4^ Data from this study improves our understanding of enterococcal adaptive evolution during recalcitrant infections, serving as foundational research for more extensive studies to define specific signatures of adaptation and correlation with clinical outcomes.

## NOTES

## Acknowledgements

The authors thank the clinical microbiology teams at each participating study site for identification and isolation of the isolates used in this study.

## Data availability

Whole-genome hybrid assemblies of isolates used in the study are available with publication on the National Institutes of Health (NIH) National Center for Biotechnology Information (NCBI) under BioProject ID PRJNA118579 and the accession numbers listed in **Supplementary Data 1.**

## Authors’ contributions

All authors contributed to the study conception and design. BMH and CAA contributed equally to this work. SRS was responsible for data cleaning, isolate sequencing, analysis, and drafting of the manuscript. PVS, MGS, AMD, DT, LB, GS, JK, VW, RC, KN, and AM were responsible for data collection. AQD, HSS, and AED were responsible for isolate processing and sequencing. All authors reviewed and approved the final version of the manuscript.

## Supplementary Material

### Supplementary Data 1

This appendix contains all isolate metadata, including BioSample accession numbers.

### Supplementary Data 2

**T1.** Genes present in *E. faecium* persistent index isolates that were not present in the comparison group isolates.

**T2.** Scoary output of *E. faecium* persistent and comparison group isolates.

**T3.** Genes present in *E. faecalis* persistent index isolates that were not present in the comparison group isolates.

**T4.** Scoary output of *E. faecalis* persistent and comparison group isolates.

**T5.** Genes gained in *E. faecium* persistent isolate pairs (index and persistent time point).

**T6.** Genes gained in *E. faecalis* persistent isolate pairs (index and persistent time point).

**T7.** Genes lost in *E. faecalis* persistent isolate pairs (index and persistent time point).

**T8.** Genes lost in *E. faecium* persistent isolate pairs (index and persistent time point).

**T9.** Genes gained in *E. faecium* recurrent isolate pairs (index and recurrent time point).

**T10.** Genes lost in *E. faecium* recurrent isolate pairs (index and recurrent time point).

**T11.** Catalogue of chromosomal structural variations in *E. faecium* and *E. faecalis* persistent isolate pairs.

**T12.** Single-nucleotide polymorphisms (SNPs) in *E. faecium* persistent isolate pairs.

**T13.** Single-nucleotide polymorphisms (SNPs) in *E. faecium* recurrent isolate pairs.

**T14.** Single-nucleotide polymorphisms (SNPs) in *E. faecalis* persistent isolate pairs.

### Isolate processing and sequencing

Isolates were obtained from each participating institution’s clinical microbiology laboratory and shipped to the central research laboratory in Houston, TX for processing and storage. For whole-genome sequencing, isolates were grown on Brain Heart Infusion (BHI) agar with or without 4 µg/mL of vancomycin depending on the reported clinical vancomycin susceptibility of the isolate. Isolates were then grown for 6-8 hours at 37°C in BHI infusion broth with or without vancomycin, then pelleted and stored at −80°C until genomic DNA could be extracted. Genomic DNA was isolated using the QIAGEN DNeasy Blood and Tissue kit (#69506), and quality and concentration were assessed with the BioTek Take3 Trio microvolume plate (Agilent Technologies) and Qubit 4 Fluorometer (Invitrogen), respectively. Library preparation for Illumina short-read sequencing was performed using the Nextera DNA Flex kit (#20018705) with IDT for Illumina DNA/RNA UD Indexes Sets A-D (#20027213-216). Isolates were sequenced with either an Illumina MiSeq, HiSeq 4000, or NextSeq 2000 using 2 x 150 bp read lengths. Library preparation for long-read whole-genome sequencing was performed with either the Oxford Nanopore Technologies Rapid Barcoding Kit (SQK-RBK110.96) or the Ligation Sequencing Kit (SQK-LSK109). Libraries were run using R9.4.1 flow cells on the GridION sequencing platform.

### Genomic data processing and assembly

Raw short-read genomic data was demultiplexed and converted to fastq files using bcl2fastq.^1^ The custom pipeline used for quality control, assembly, and initial analyses of short-read data (https://github.com/Hanson-Lab/CRACKLEII_pipeline) includes raspberry-v0.3^2^ for examination of short reads for quality and adaptor contamination, and bases under a Phred quality score of 30, along with adaptors, were trimmed using Trimmomatic-v0.39^3^. Next, assemblies were created with SPAdes-v3.15.3^4^ and qualitatively examined with QUAST-v5.0.2^5^. Lastly, *in silico* multi-locus sequence typing (MLST) was performed with mlst-v2.19.0,^6,7^ and assemblies were annotated for downstream use with Prokka-v1.14.5^8^

Long-read genomic sequencing data was basecalled with Guppy-v4.5.2,^9^ and adaptors and reads <500 bases were removed using Trimmomatic.^3^ A custom, in-house python script^10^ was used to create highly contiguous, circular hybrid assemblies utilizing both short- and long-read sequence data. Briefly, this script uses Flye-v2.9 for the creation of *de novo* long-read assemblies, which are then circularized with berokka-v0.2.^12^ Assemblies are oriented to *dnaA*, and redundant and self-contained contigs are removed using circlator’s ‘fixstart’ and ‘clean’ commands, respectively.^13^ Long reads undergo iterative polishing with Racon-v1.4.21^14^ and medaka-v0.8.1,^15^ followed by four rounds of short-read polishing with Racon.^14^ Quality control is performed using Snippy-v4.6.0^16^ to identify erroneous SNPs from the long-read assemblies that are corrected using the highly accurate short-read data. Short- and long-read pileups using minimap2^17^ is then used to assess read coverage of assemblies. Final hybrid assemblies were annotated with Prokka-v1.14.5,^8^ and antimicrobial resistance gene and plasmid content were assessed with the AMRFinderPlus^18^ and PlasmidFinder^19^ databases, respectively.

### Analysis of clinical data

Composite clinical measures that include Charlson Comorbidity Index^20^ and Pitt Bacteremia Score^21^ were computed based on available clinical data. Empiric antibiotic therapy was defined as any anti-enterococcal therapy initiated before culture results were finalized. Definitive therapy was defined as anti-enterococcal therapy initiated within 24 hours after the index culture results were finalized, or empiric therapy that was continued at least 48 hours after final culture results were reported. Definitive antibiotic therapies started within seven days of final culture date and with at least 48 hours of overlapping administration were classified as “combination definitive therapy.” Appropriate therapy was defined as receipt of ≥ 1 day of anti-enterococcal antibiotics with *in vitro* activity as a single agent (i.e., linezolid monotherapy against linezolid-susceptible *E. faecalis*) or as part of a combination therapy (i.e., daptomycin plus ampicillin against ampicillin-resistant *E. faecium*). Inappropriate therapy was defined as receipt of a single antibiotic without evidence of *in vitro* activity (i.e., ampicillin monotherapy against ampicillin-resistant *E. faecium*). To evaluate group-level differences in the persistent/recurrent vs. comparison groups (binary outcome), univariable statistics for continuous variables were computed using non-parametric Wilcoxon rank sum tests. For categorical data, either Pearson’s chi-square or Fisher’s exact tests were used depending on sparsity of the data. Variables that had a significance level p ≤0.1 were included in a generalized linear mixed model (GLMM) using the glmnet package ^22^ with institution of admission included as a random effect to account for potential institution-specific confounding. As antibiotic therapy was assessed separately by species, these variables were not included in the GLMM.

### Analysis of genomic features

#### Characterization of accessory genes in persistent and comparison isolates

Comparisons between persistent and comparison group index isolates, as well as between persistent index isolates and persistent time point isolates were performed separately for *E. faecium* and *E. faecalis*. Genes were considered accessory if they were present in 95% or less of isolates in the cohorts being compared. Because closed genomes were generated for all isolates, contigs other than the chromosome determined to be circular by Flye and by manual inspection using Integrative Genomics Viewer (IGV) v2.13.2^23^ were considered plasmid sequences, even if no *rep* gene was identified using the PlasmidFinder database.^19^ Antimicrobial resistance genes and mutations were identified using AMRFinderPlus v3.11.8 with the respective species-specific database and a minimum identity and coverage cutoff of 0.8.

Subtype/strain determination of sequential isolates was performed using Snippy-v4.6.0.^16^ Isolates were considered the same subtype, or strain, if the pairwise whole-genome pairwise SNP distance was <20 SNPs. This threshold is based on Gouliouris et. al,^24^ who found that isolates containing less than 20 SNPs exhibited limited recombination and were of the same ST, which was consistent with our findings in our own cohort.

Genes were mapped to functional pathways via orthology using the Kyoto Encyclopedia of Genes and Genomes (KEGG) database implemented with eggNOG-mapper-v2.1.12.^25^ Average proportions of accessory genes assigned to each clusters of orthologous group (COG) functional categories were computed, and the relative frequencies of these COG groups were compared across persistent and comparison isolate groups, as well as persistent infection pairs. Scoary^26^ was used to identify genes that were significantly associated with persistence using a naive p-value cutoff of 0.05.

#### Functional characterization of genes with mutations arising during persistent bacteremia

Mutations were catalogued in all genes in persistent time point isolates using Snippy-v4.6.0,^16^ with each index isolate serving as the patient-specific reference. Mutation-containing genes were functionally annotated using eggNOG as described above, and the pairwise relative frequencies of identified COG groups were calculated for each infection for both *E. faecium* and *E. faecalis* persistent infection pairs.

#### Characterization of chromosomal structural variants

Individual chromosomes for each persistent infection pair were compared using NucDiff v2.0.3^27^ to obtain the location and type of longitudinal chromosomal structural variations (SV) present in persistent bacteremia. The index blood culture isolate served as the reference sequence for each respective infection pair. Due to the high level of enterococcal genomic plasticity and its proclivity for recombination, SV >1000bp were considered for further analyses. All chromosomal SVs were visualized using IGV.^23^

### Supplementary Tables

**Table S1.**
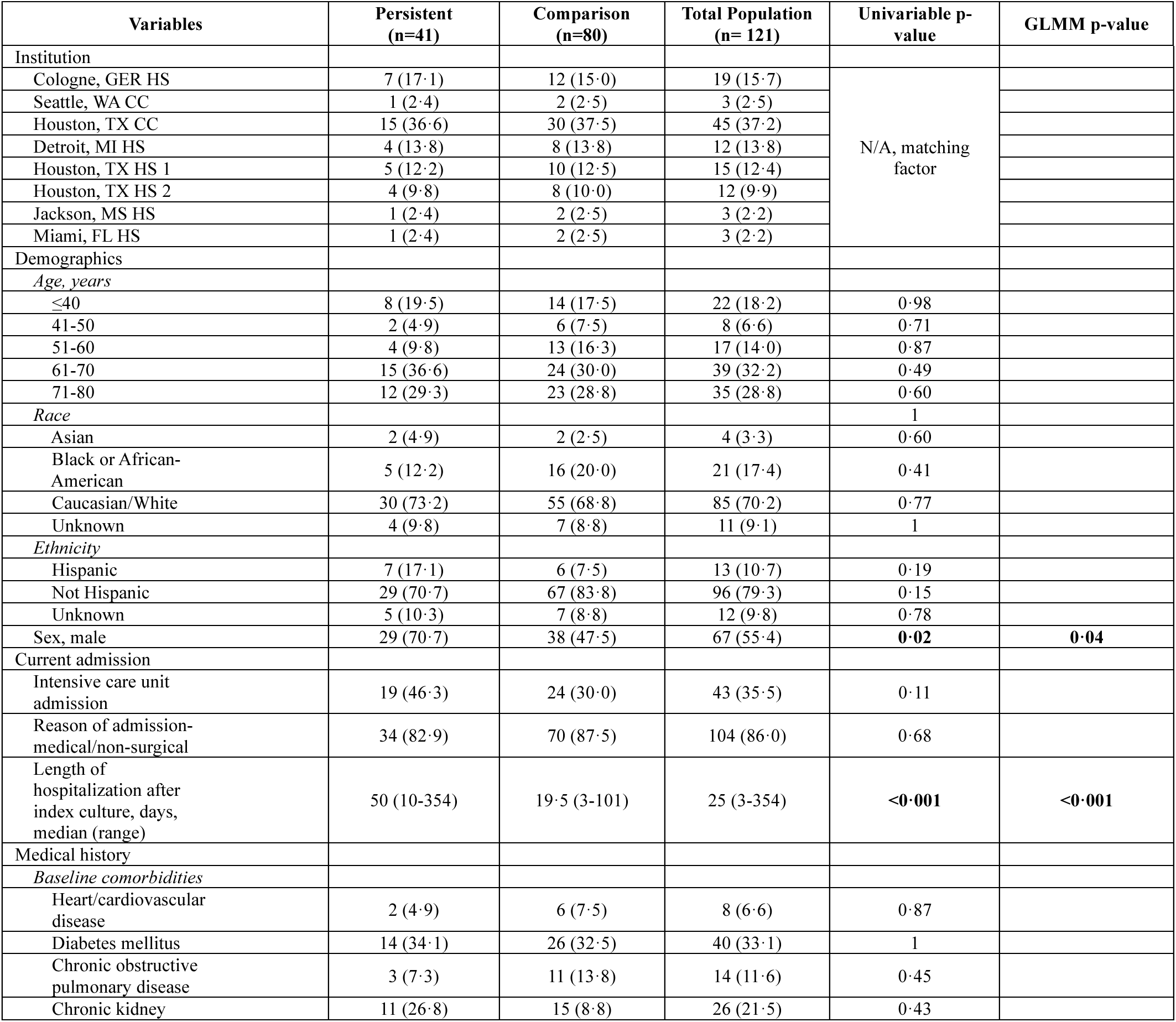
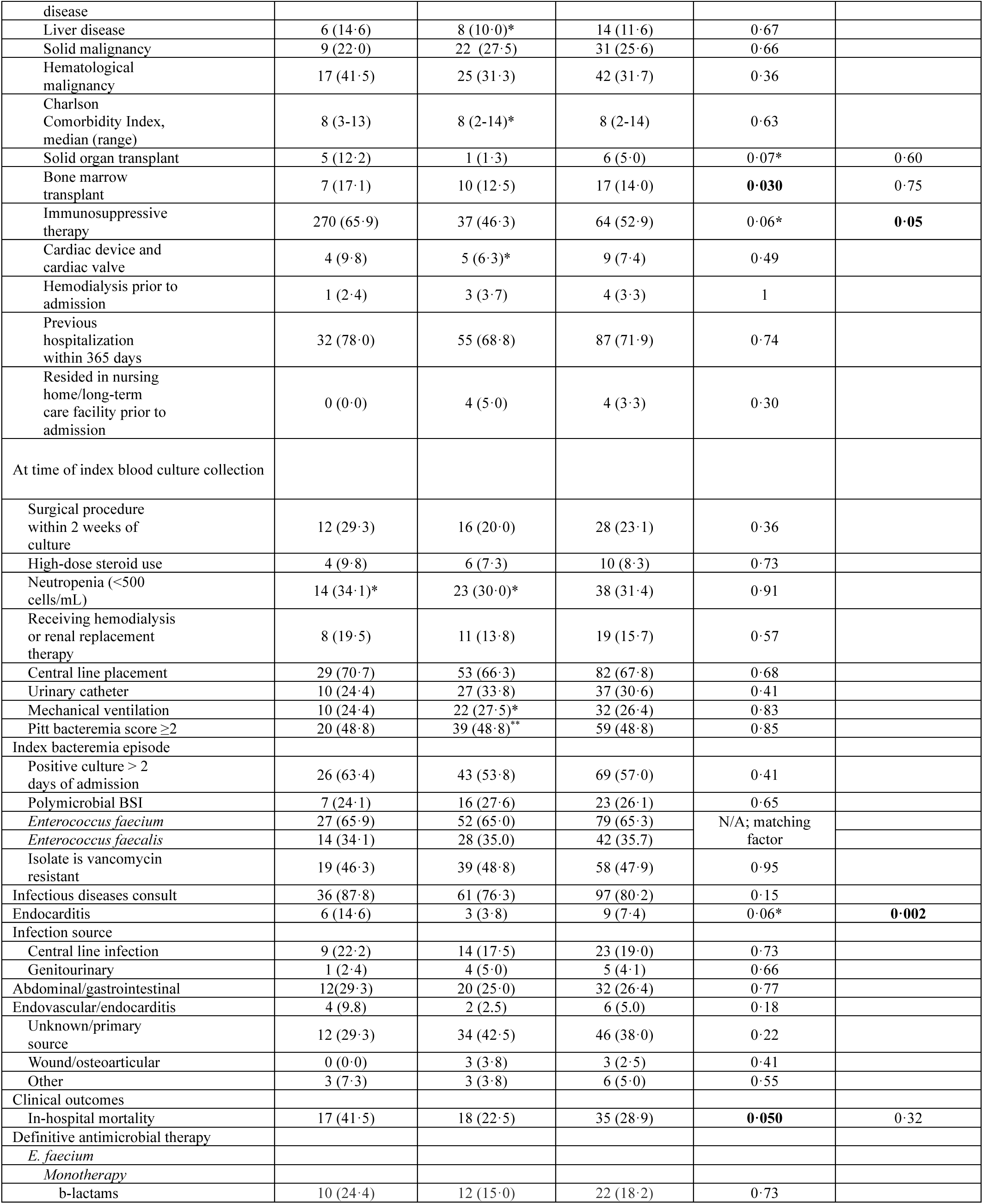
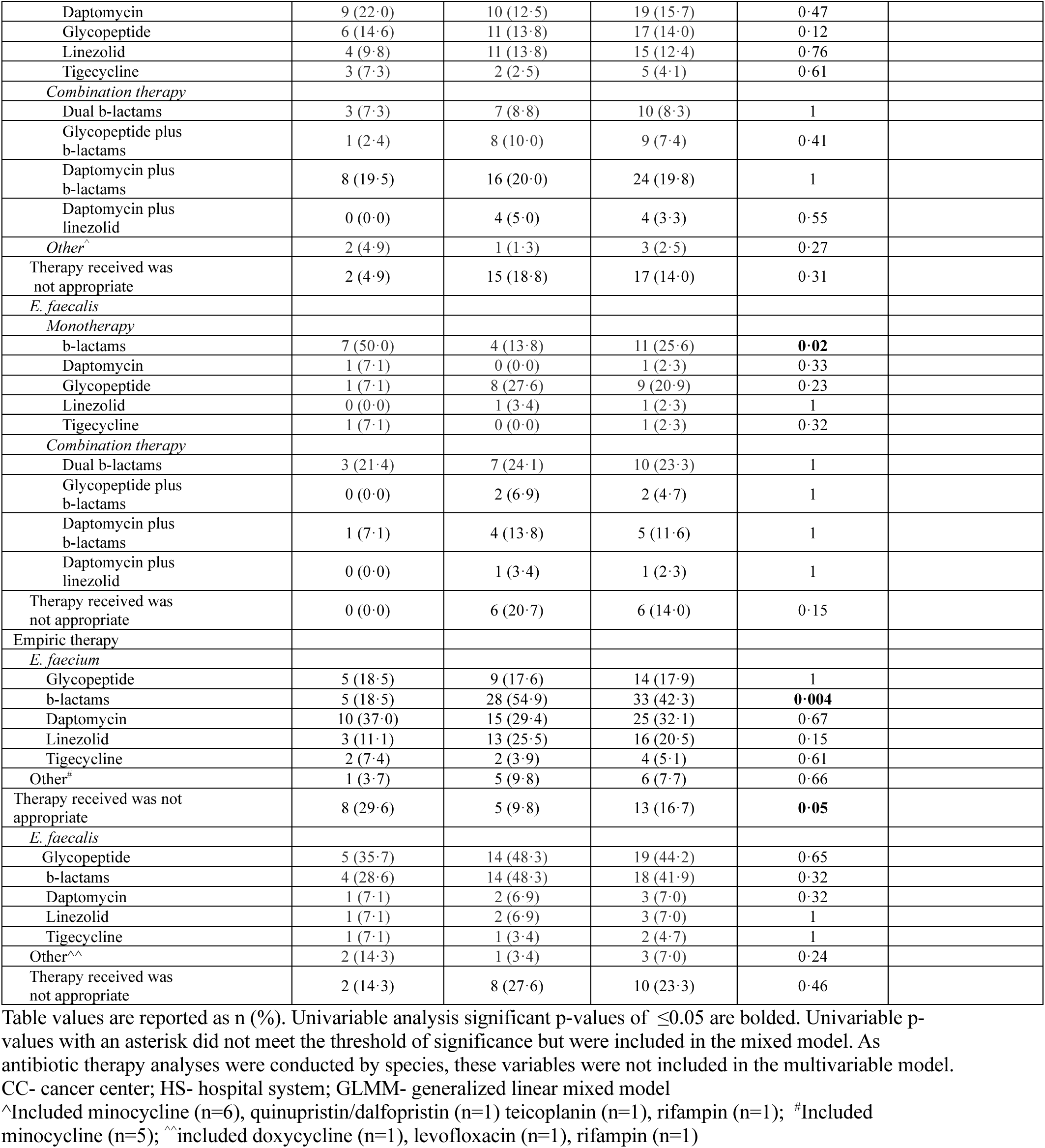
Clinical characteristics of patients with recalcitrant enterococcal bacteremia and comparison patients.

**Table S2.**
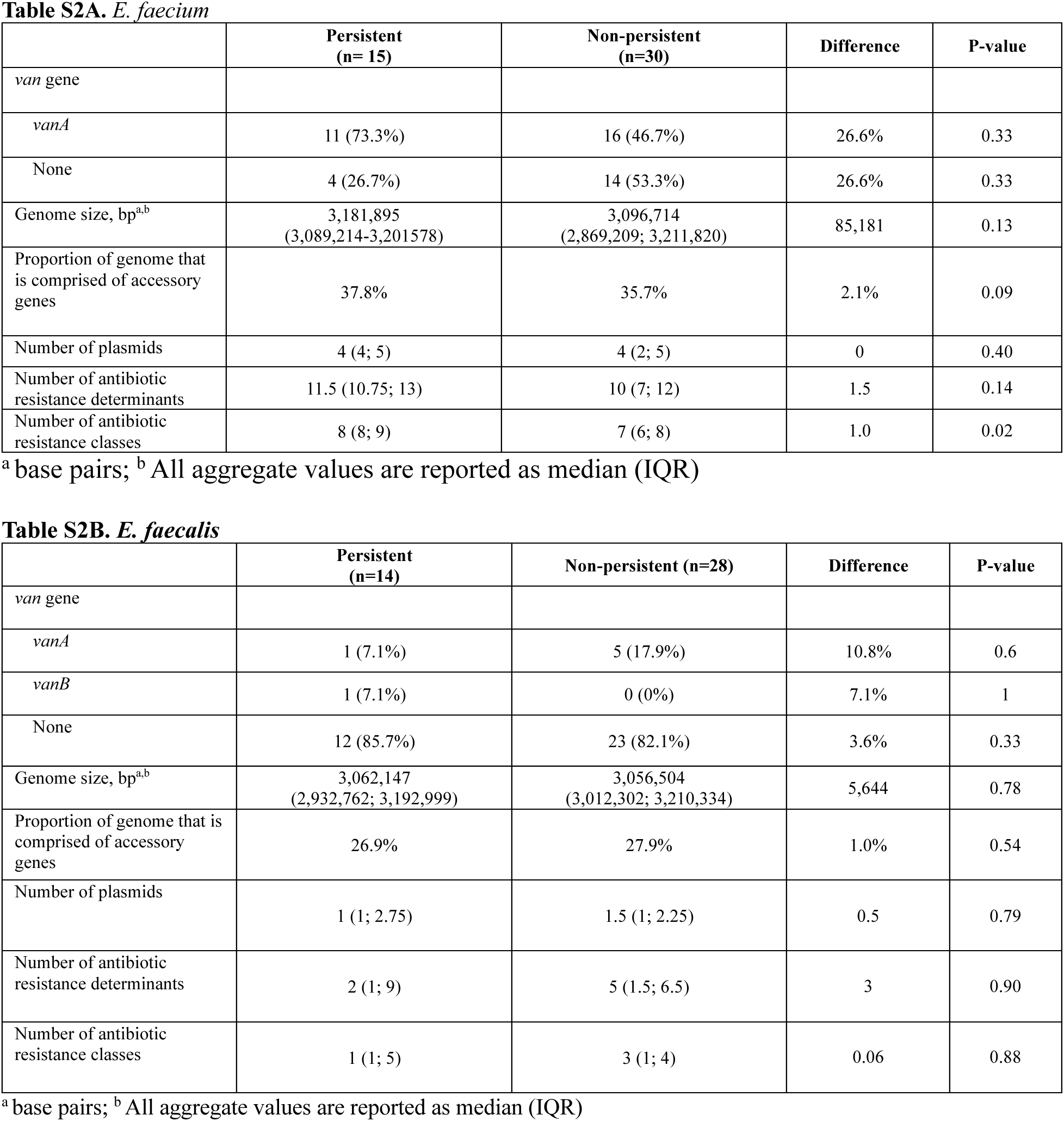
Descriptive genomic comparisons between persistent and non-persistent enterococcal infections.

**Table S3.**
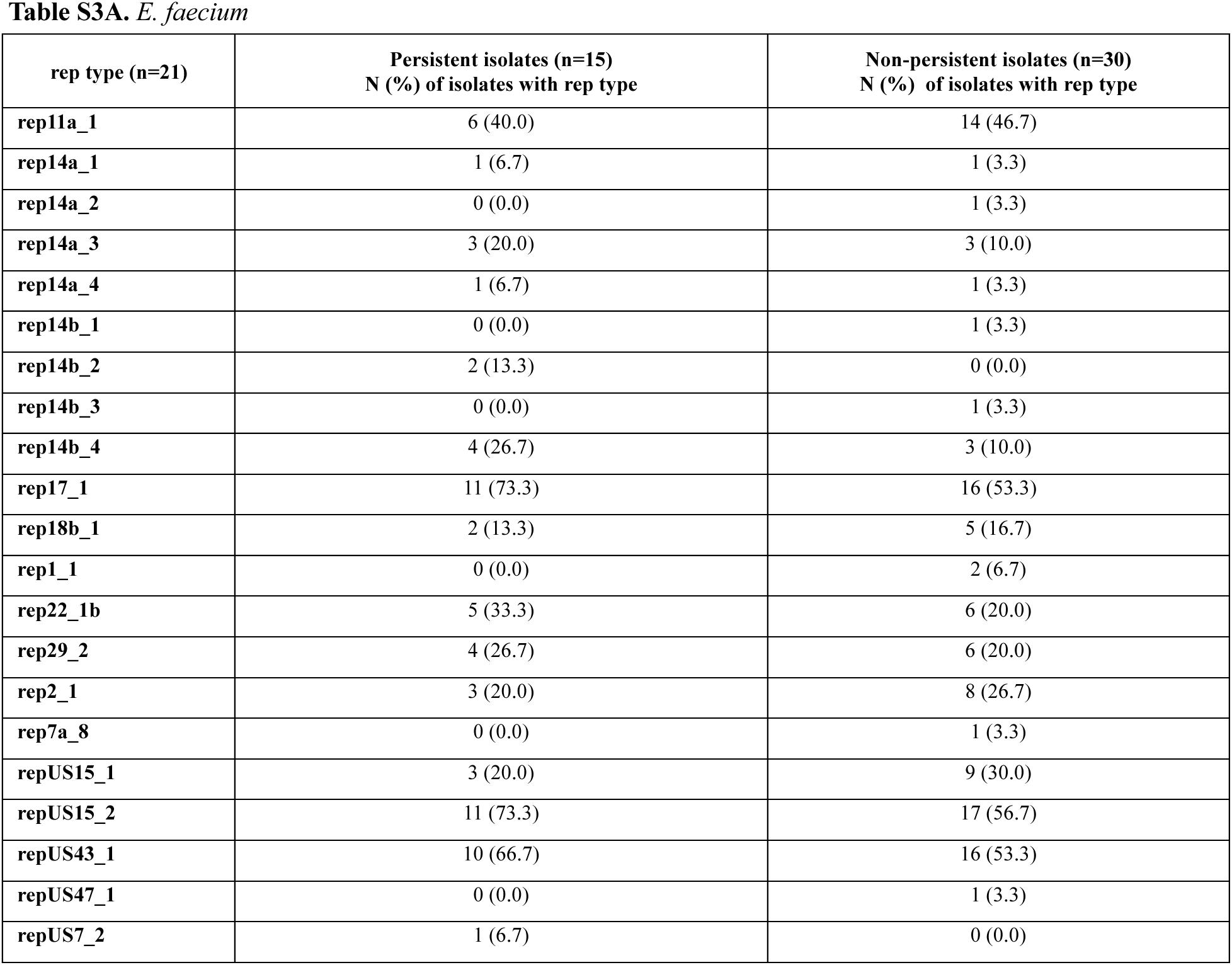
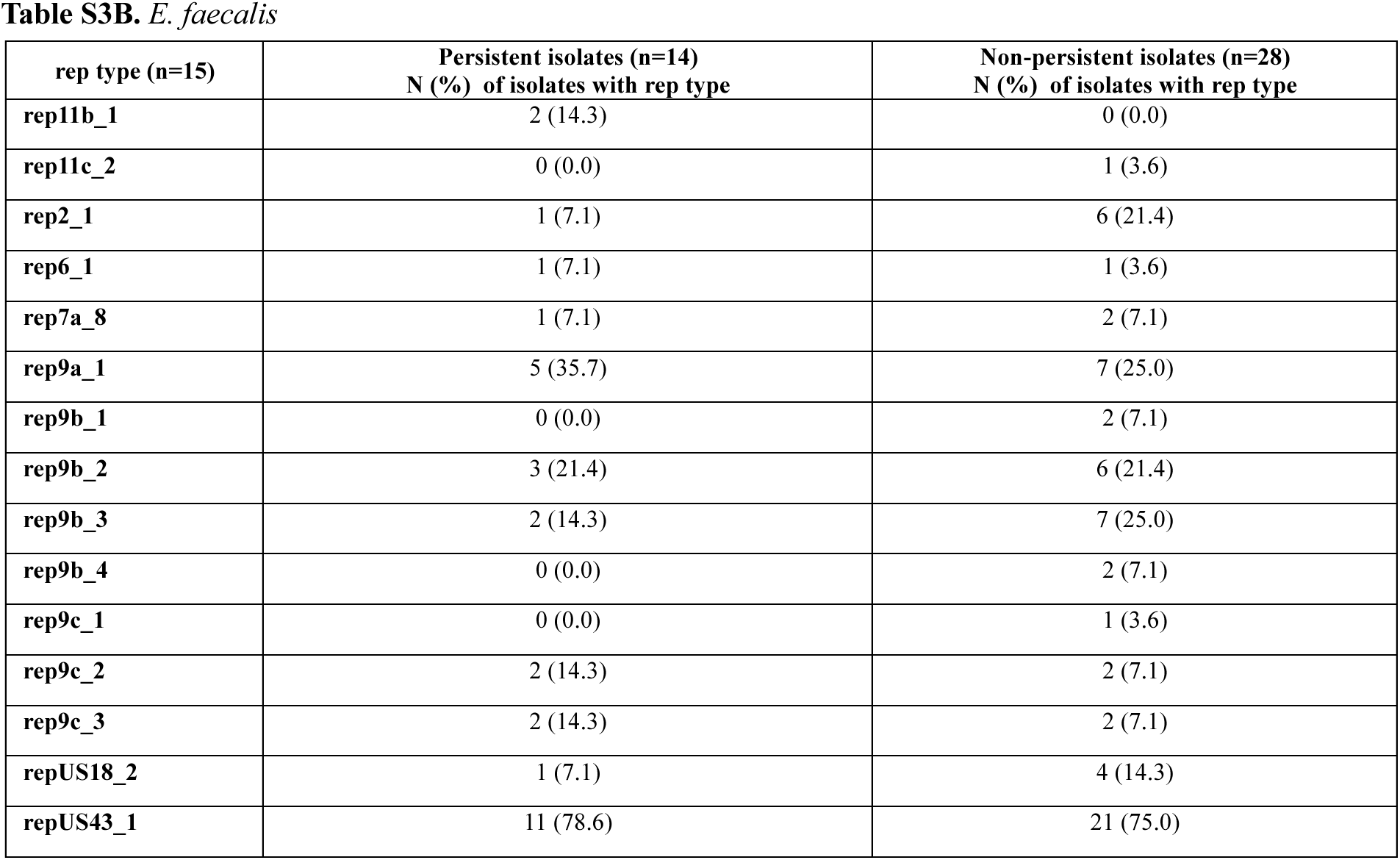
Presence of rep types in persistent vs. non-persistent isolates.

**Table S4.**
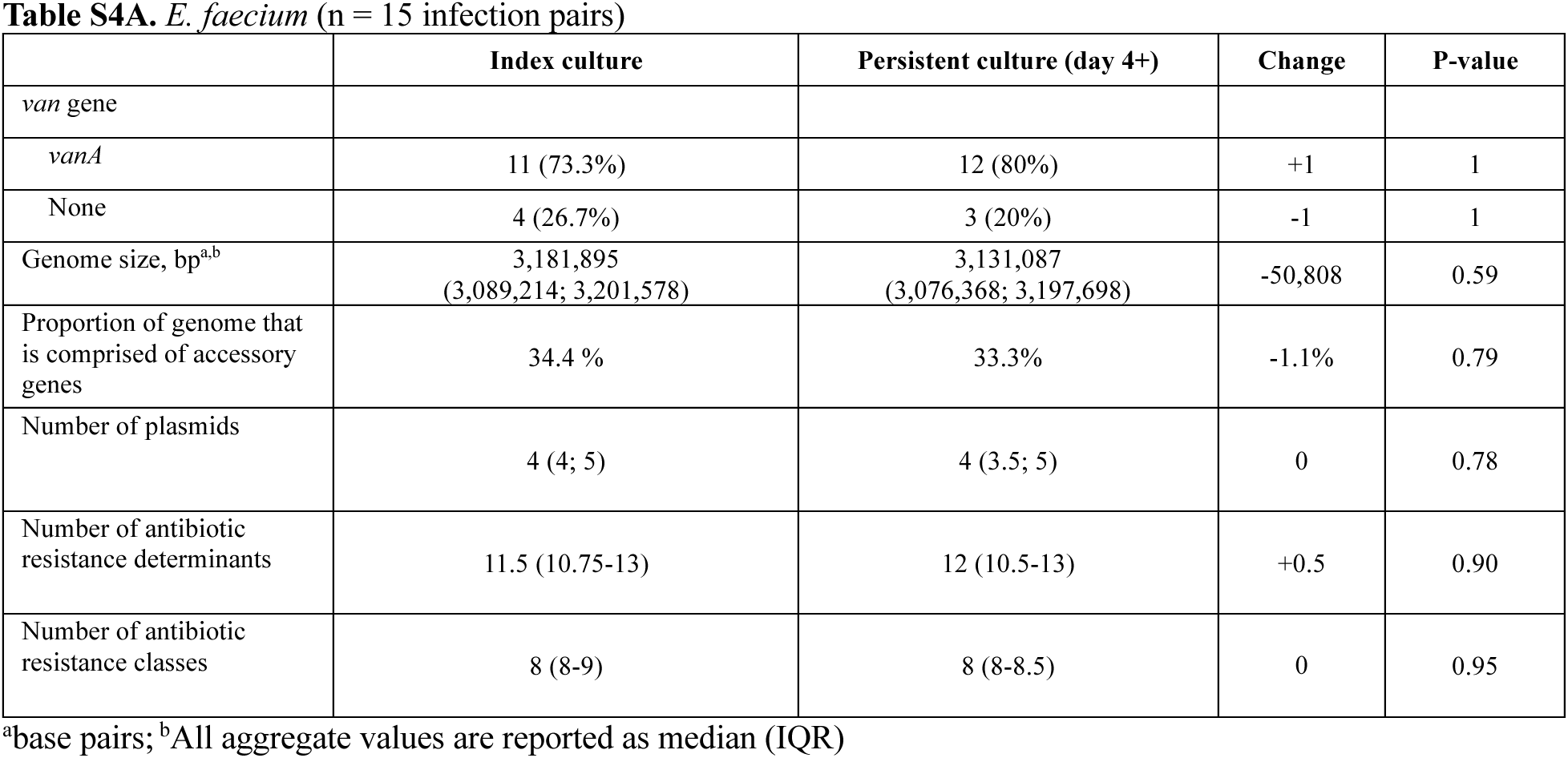
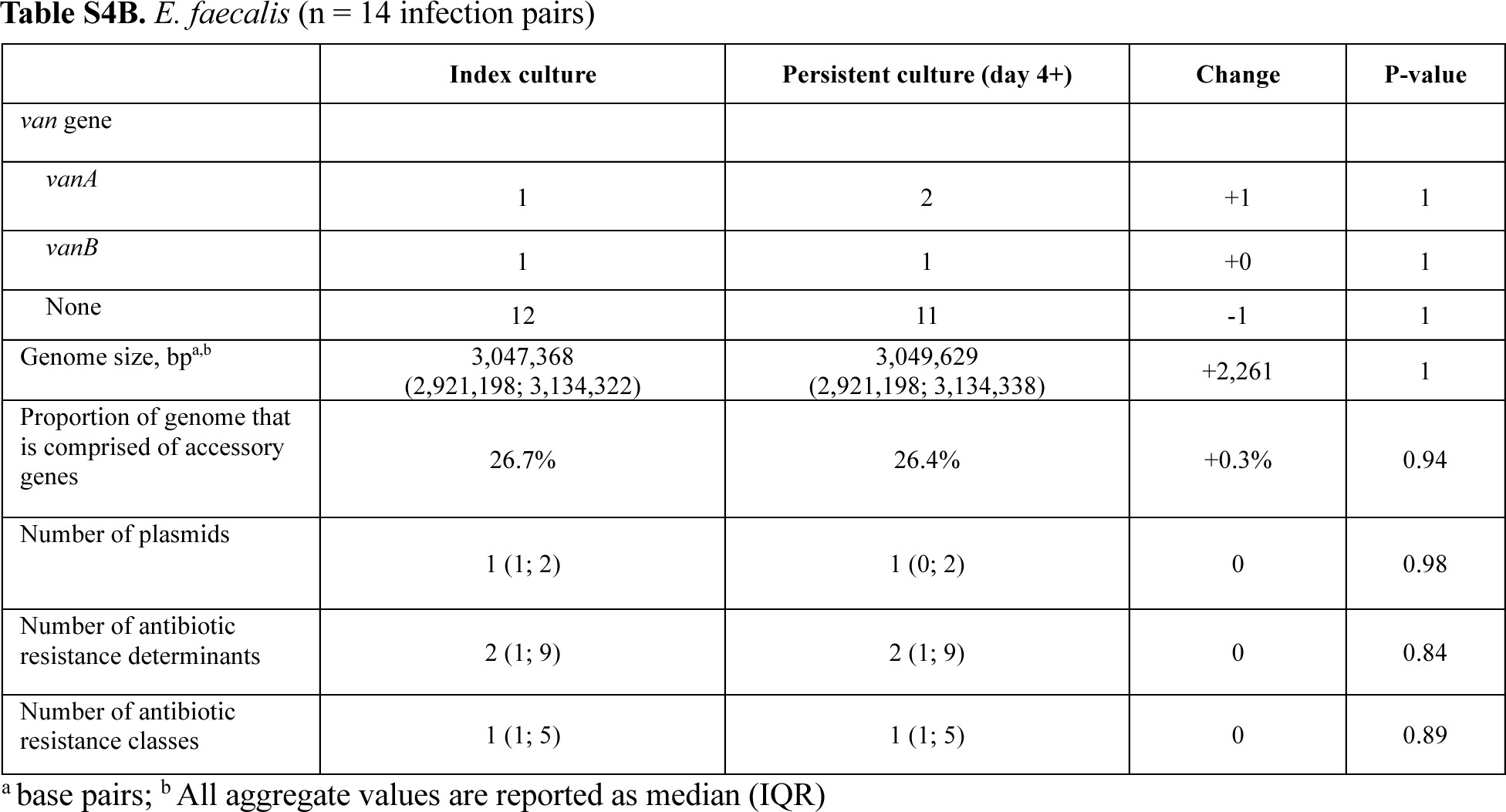
Descriptive genomic comparisons between index isolates of persistent infections and isolates at day 4+ (persistent time point).

**Table S5.**
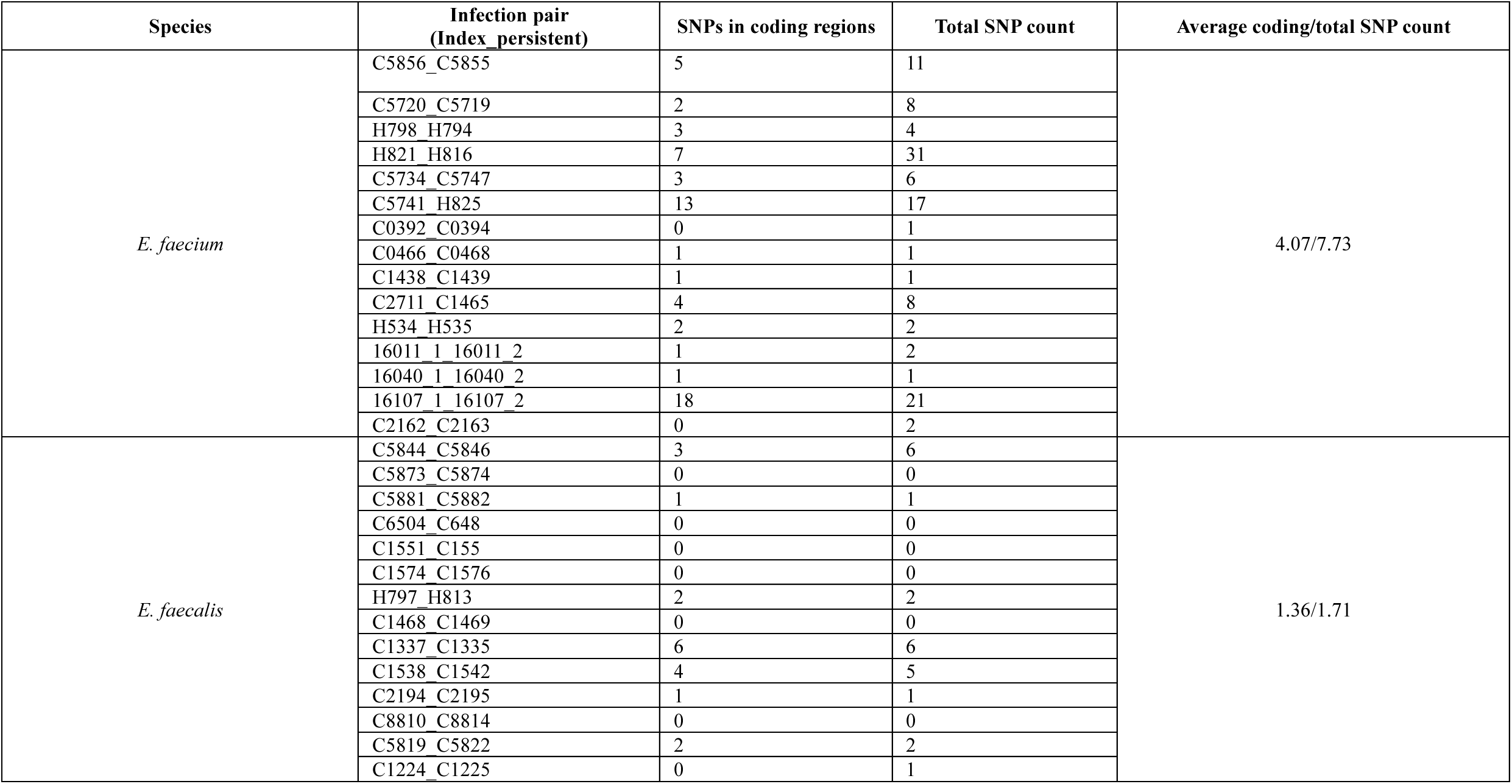
SNP counts between index persistent isolates and persistent time point isolates.

### Supplementary Figures

**Figure S1.**
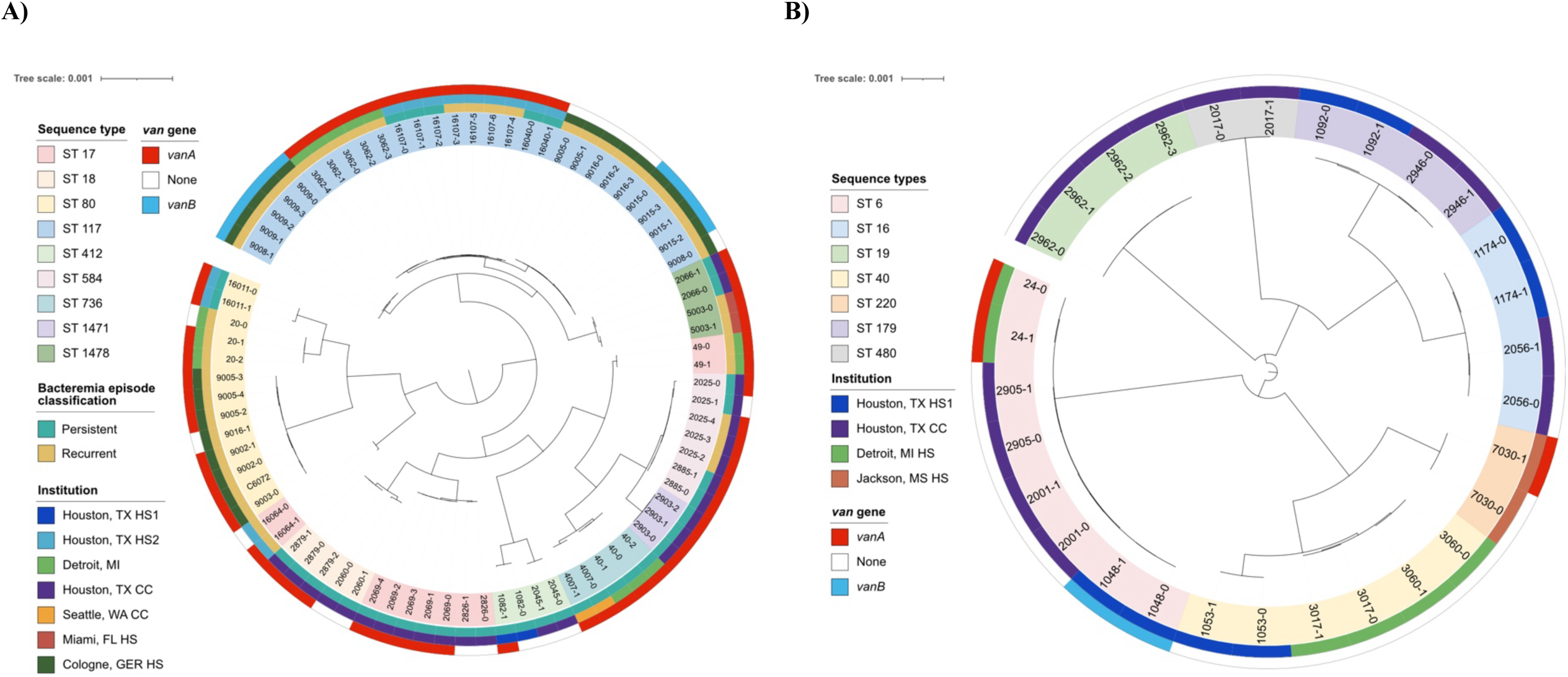
Population structure of enterococcal isolates causing persistent bacteremia. A) *E. faecium*; B) *E. faecalis*. Midpoint-rooted, core genome maximum-likelihood phylogenetic trees of recalcitrant bacteremia isolates. The branch labels indicate the patient ID and time point; for example, 3060-0 corresponds to patient 3060-[index isolate], and 3060-1 corresponds to patient 3060-[first recalcitrant time point isolate]. Label shading corresponds to isolate sequence type, and colored rings denote bacteremia episode classification (for *E. faecium* only; there were no recurrent *E. faecalis* isolates available for sequencing), institution of isolate collection, and presence/absence of *van* genes.

**Figure S2.**
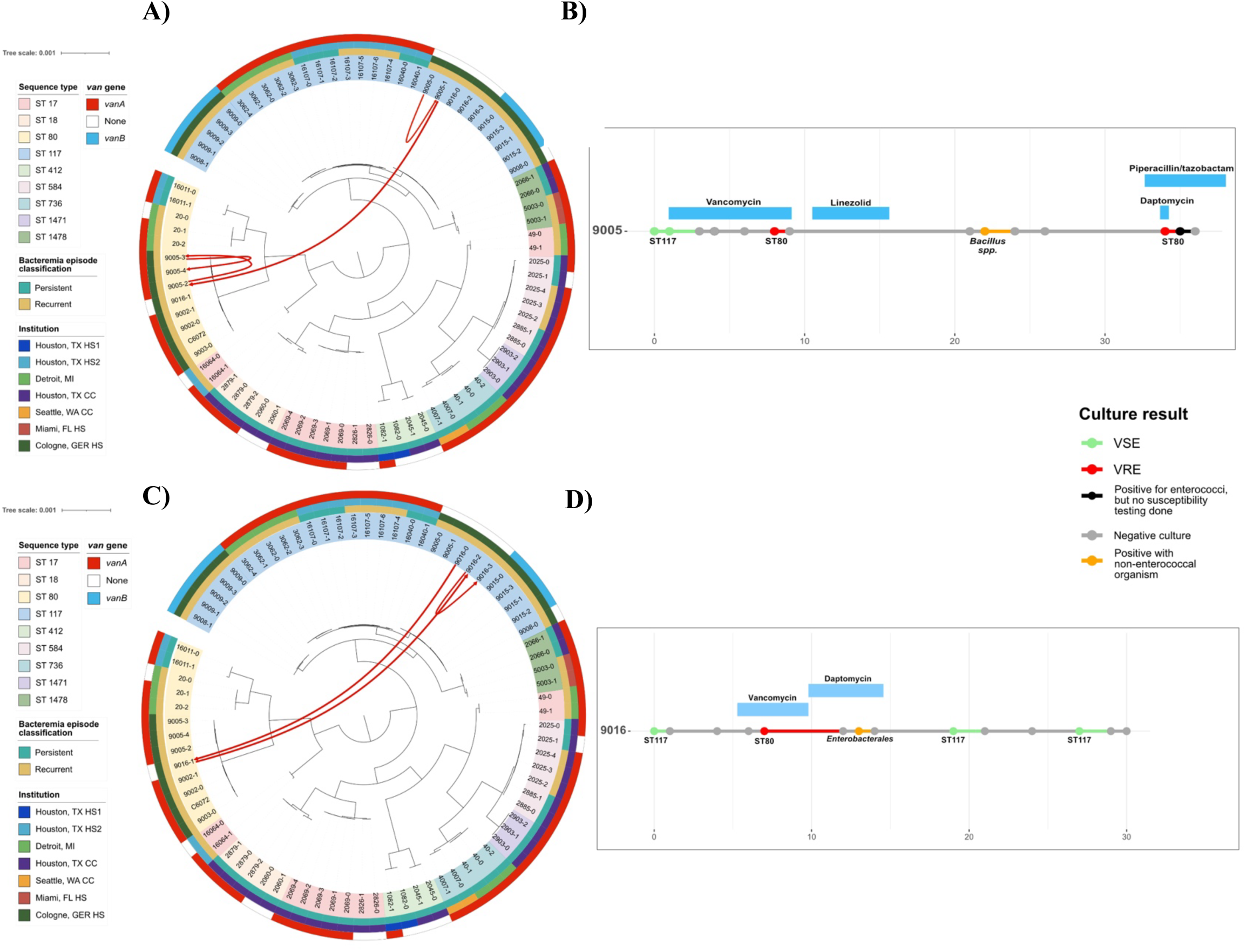
Clinical course of patients with demonstrated strain switching during recurrent bacteremia episodes. **A)** Maximum-likelihood phylogenetic tree of *E. faecium* persistent and recurrent isolate sets. Red lines correspond to blood culture isolates from patient 9005. **B)** Clinical course of patient 9005. Blue rectangles correspond to length of administration of each respective antibiotic therapy. **C)** Maximum-likelihood phylogenetic tree of *E. faecium* persistent and recurrent isolate sets. Red lines correspond to blood culture isolates from patient 9016. **D)** Clinical course of patient 9016. SE = vancomycin-susceptible enterococci; VRE = vancomycin-resistant enterococci; HS = hospital system; CC = cancer center

**Figure S3.**
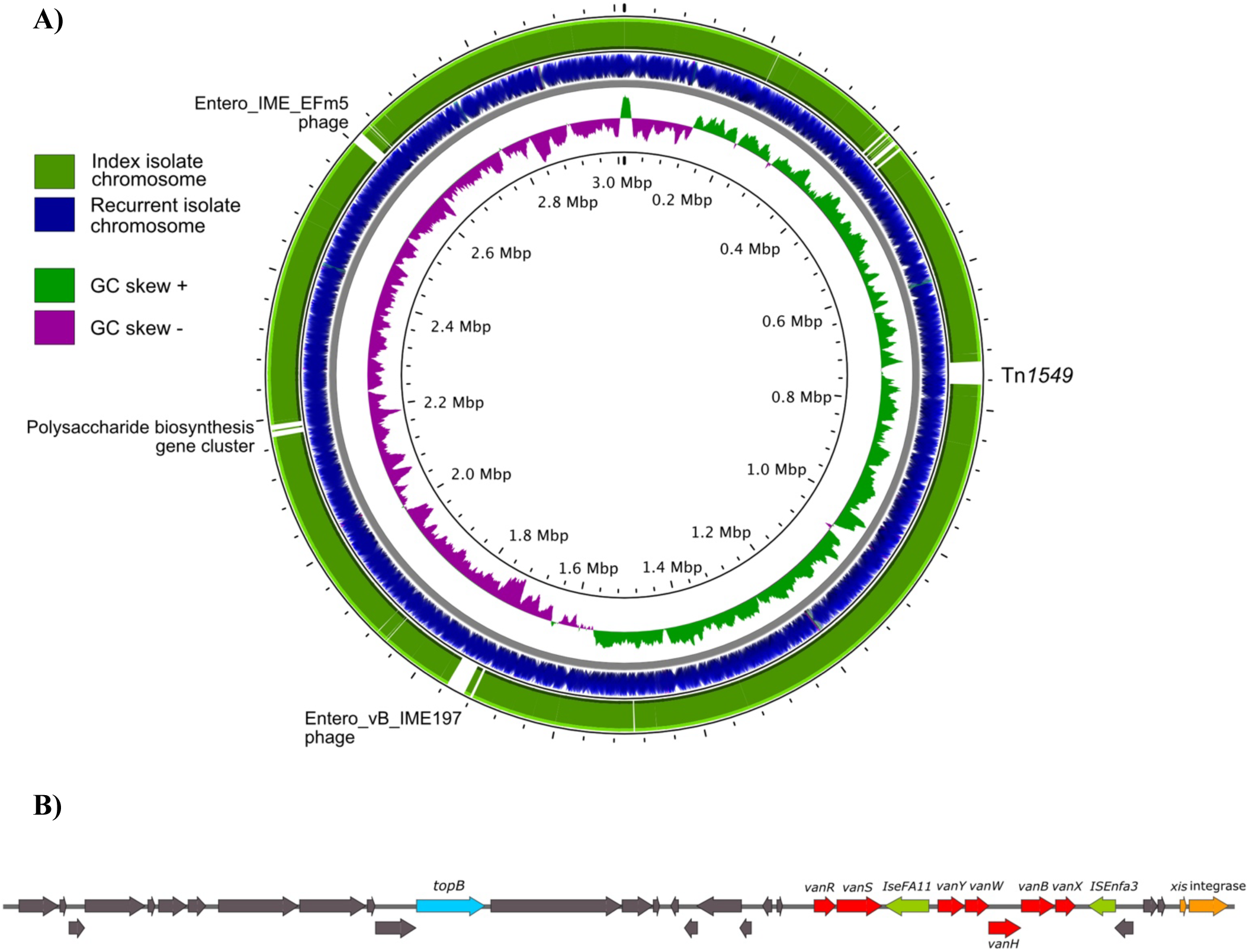
Characterization of isolates from a recurrent bacteremia episode with atypically high SNP counts **A)** Comparison of chromosomes and G/C content skew. **B)** Inserted Tn*1549* harboring *vanB* operon. The *vanB* operon components are denoted in red, hypothetical proteins in gray, insertion sequences (IS elements) in green, and excision and integration machinery in orange.

**Figure S4.**
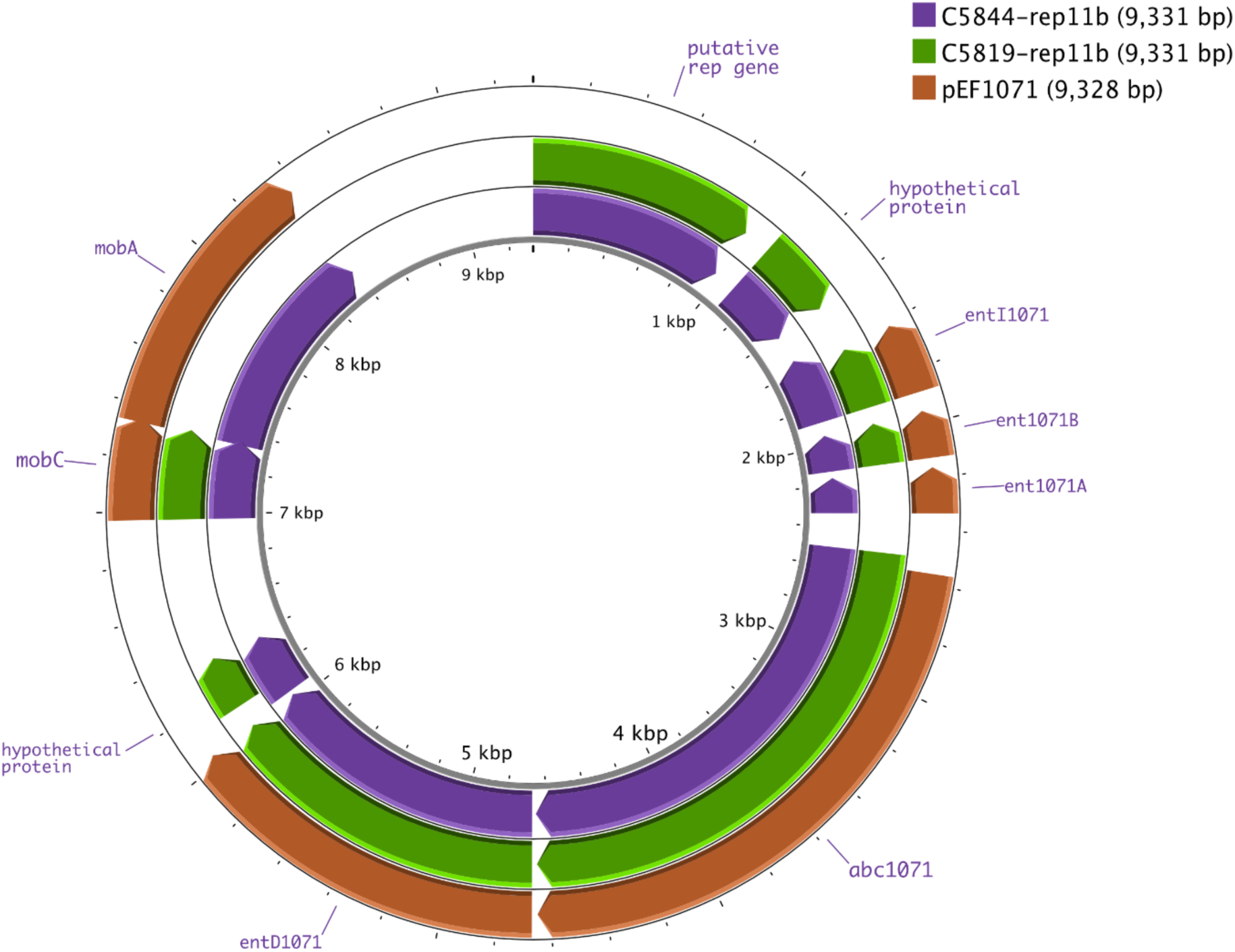
Alignment of rep11b_1-containing plasmids from persistent *E. faecalis* isolates to reference plasmid pEF1071. Purple and green circles correspond to rep11b_1-type plasmids harbored by isolates from this study. The orange ring corresponds to the reference pEF1071 sequence sequenced by Balla et al ^28^.

**Figure S5.**
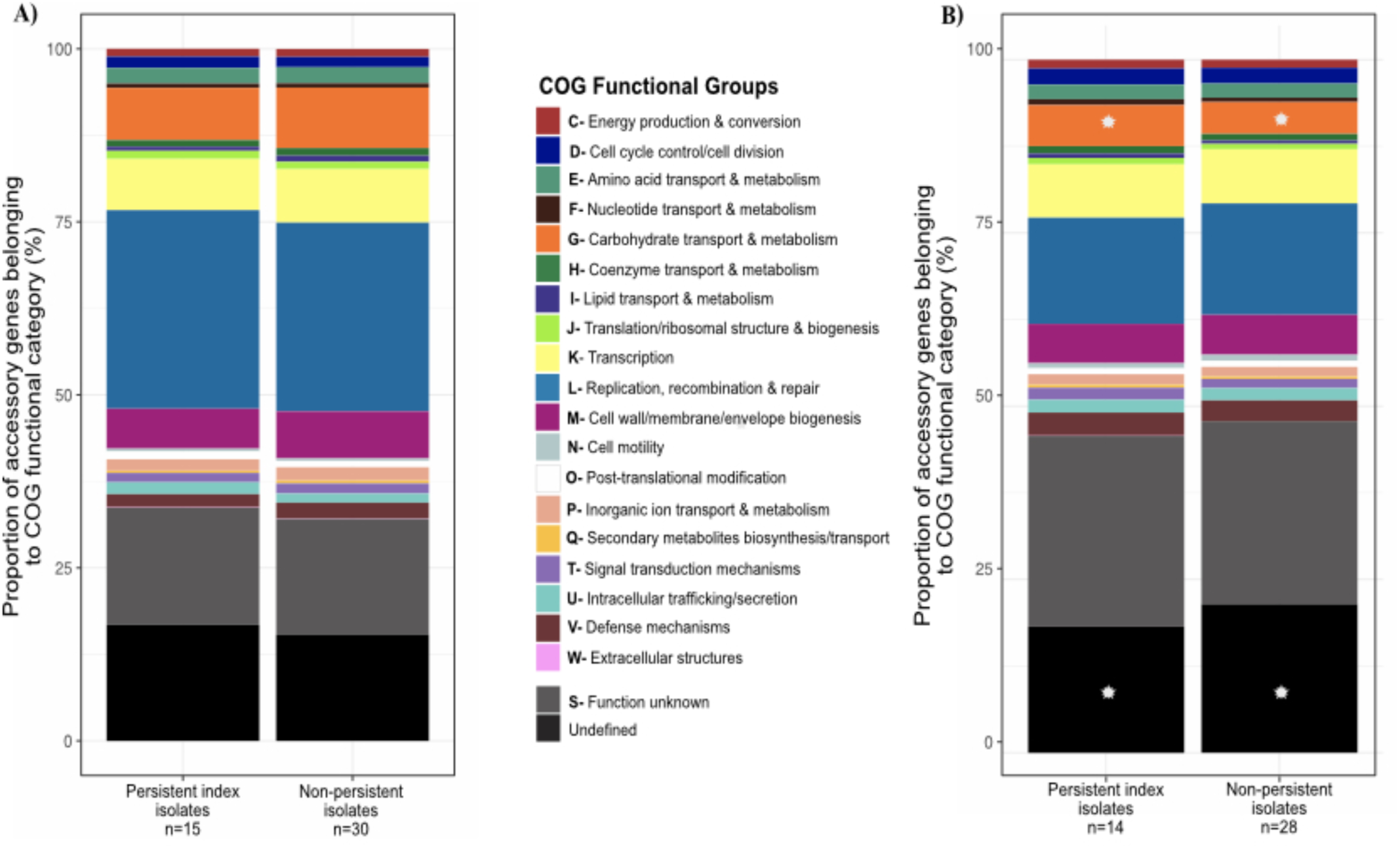
Functional characterization of accessory genes in persistent vs. non-persistent isolates. (A) *E. faecium*. (B) *E. faecalis*. These stacked bar plots indicate the average proportions of the functional classes of genes present in the accessory genomes (present in <95% of isolates) of persistent vs. non-persistent comparison isolates. A white star denotes a significant (p < 0.05) difference in proportion of genes belonging to the respective functional group between bacteremia groups.

**Figure S6.**
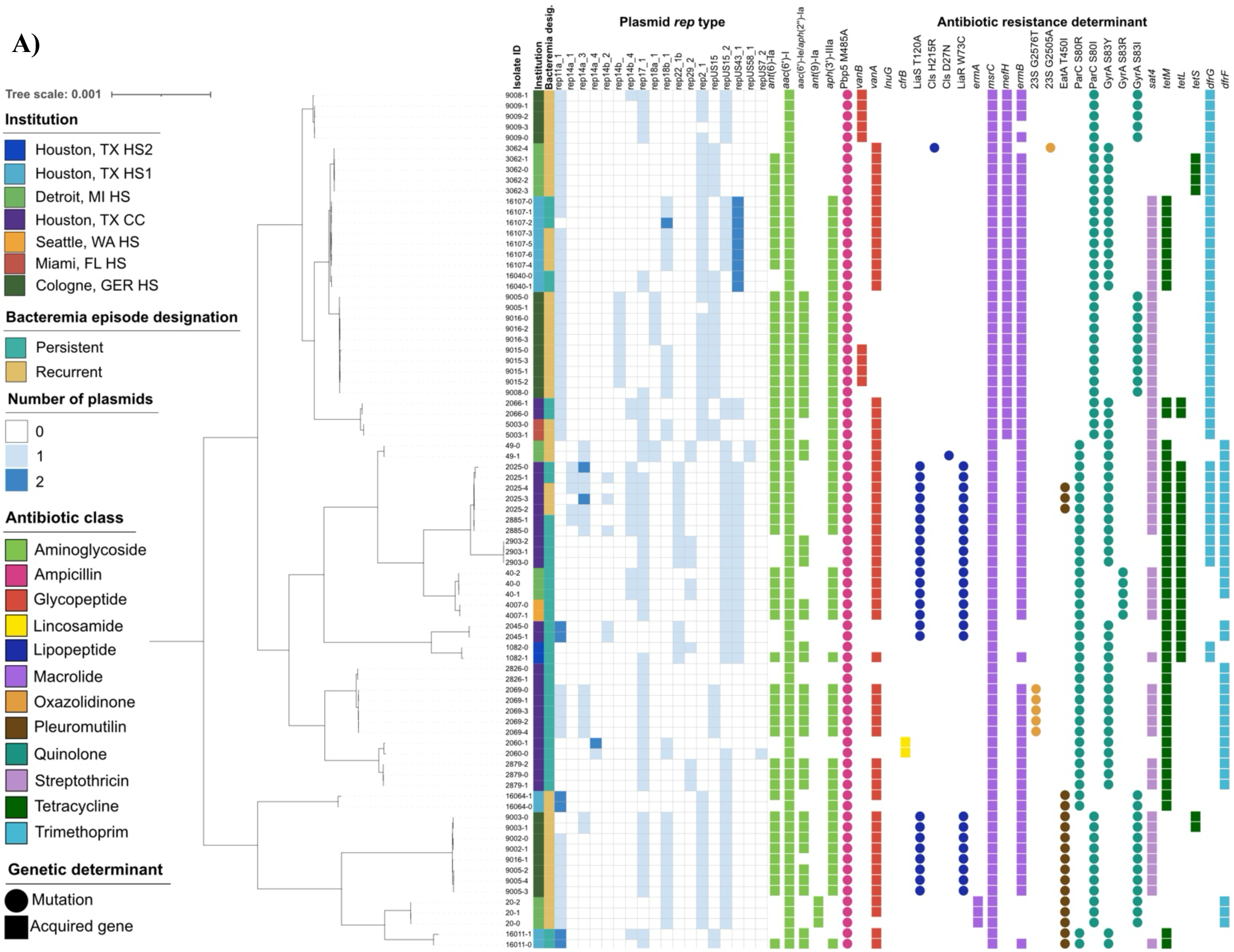
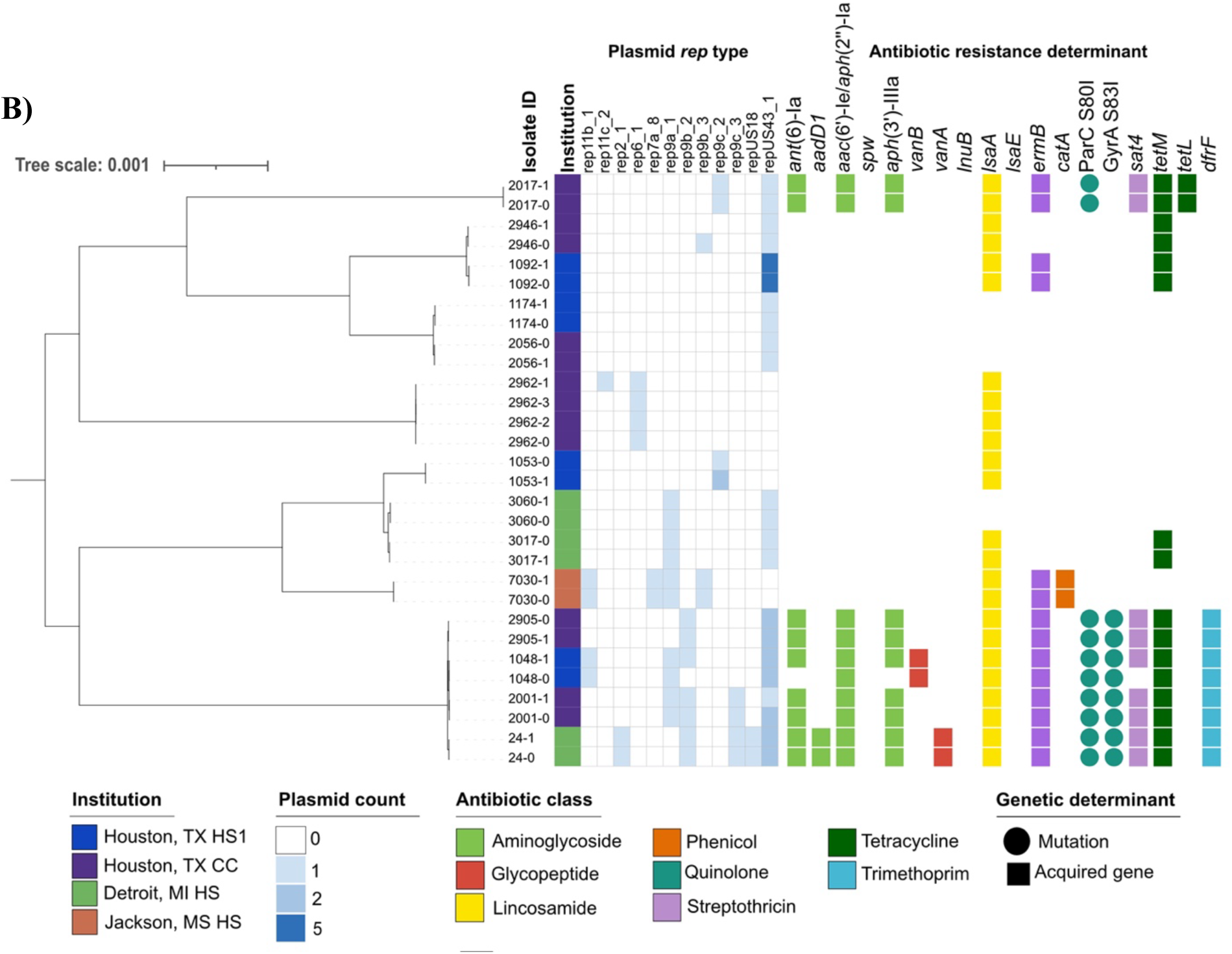
Typeable plasmid and antimicrobial resistance determinant repertoires of recalcitrant *E. faecium* and *E. faecalis* isolates A) *E. faecium*. B). *E. faecalis*. Maximum-likelihood phylogenetic trees of recalcitrant *E. faecium* and *E. faecalis* bloodstream isolates are overlaid with the typeable plasmid repertoire (left) and antimicrobial resistance determinant reservoir (right). Isolate IDs correspond to the unique combination of patient ID and blood culture time point, where “-0” corresponds to the index isolate.

**Figure S7.**
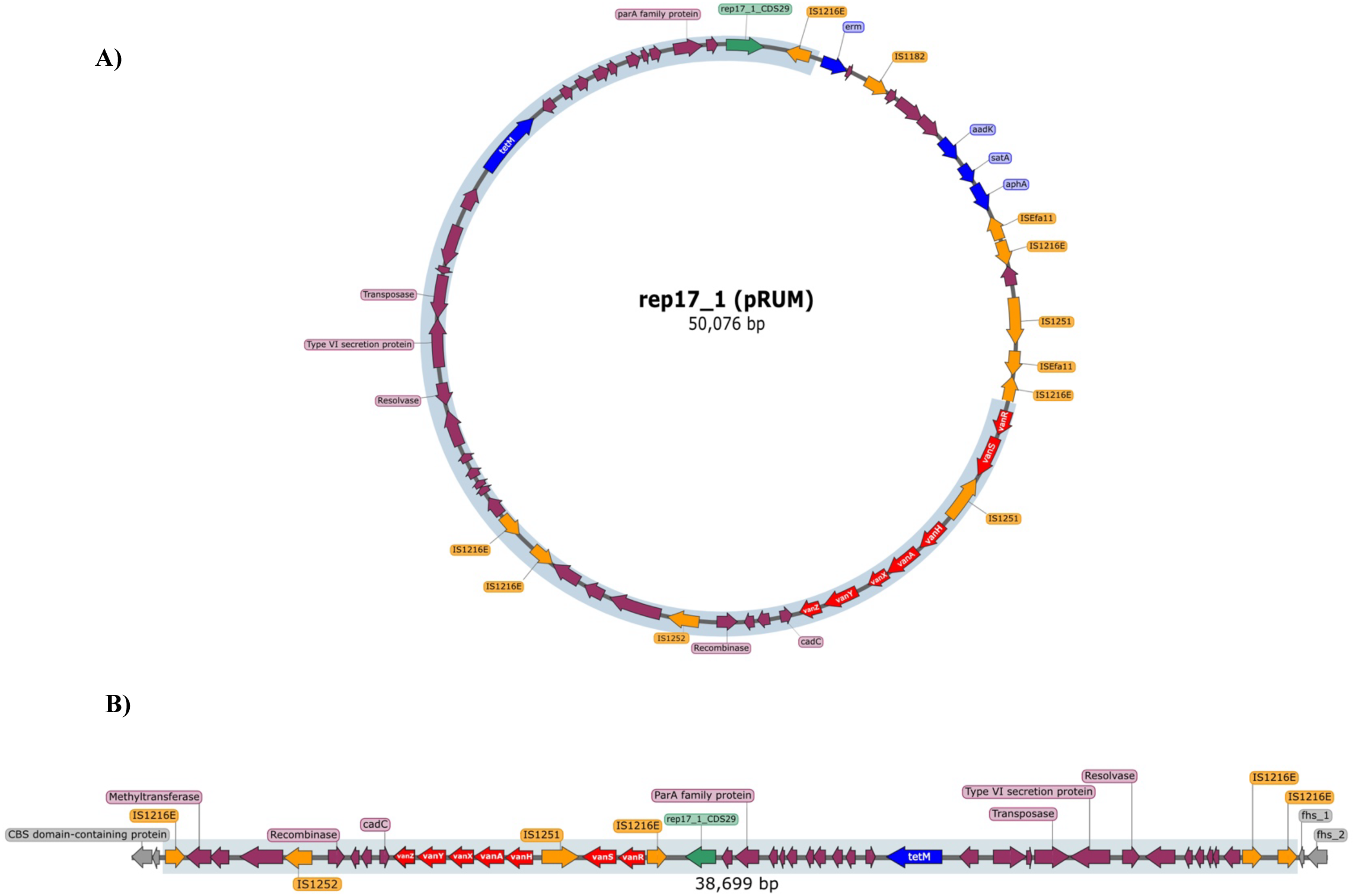
Integration of *E. faecium* index isolate plasmid harboring *vanA* operon into the chromosome of the persistent isolate. **A)** Gene map of the 50kbp pRUM plasmid harboring the *vanA* operon in the index *E. faecium* bloodstream isolate. Purple genes correspond to hypothetical proteins. The light blue shaded region corresponds to the sequence that was integrated into the chromosome of the persistent isolate. **B)** Gene map of the integration site of the index pRUM plasmid into the chromosome of the persistent isolate. Gray genes correspond to regions of the chromosome that were not part of the integration event. The blue-gray shaded region corresponds to the integrated plasmid sequence.

**Figure S8.**
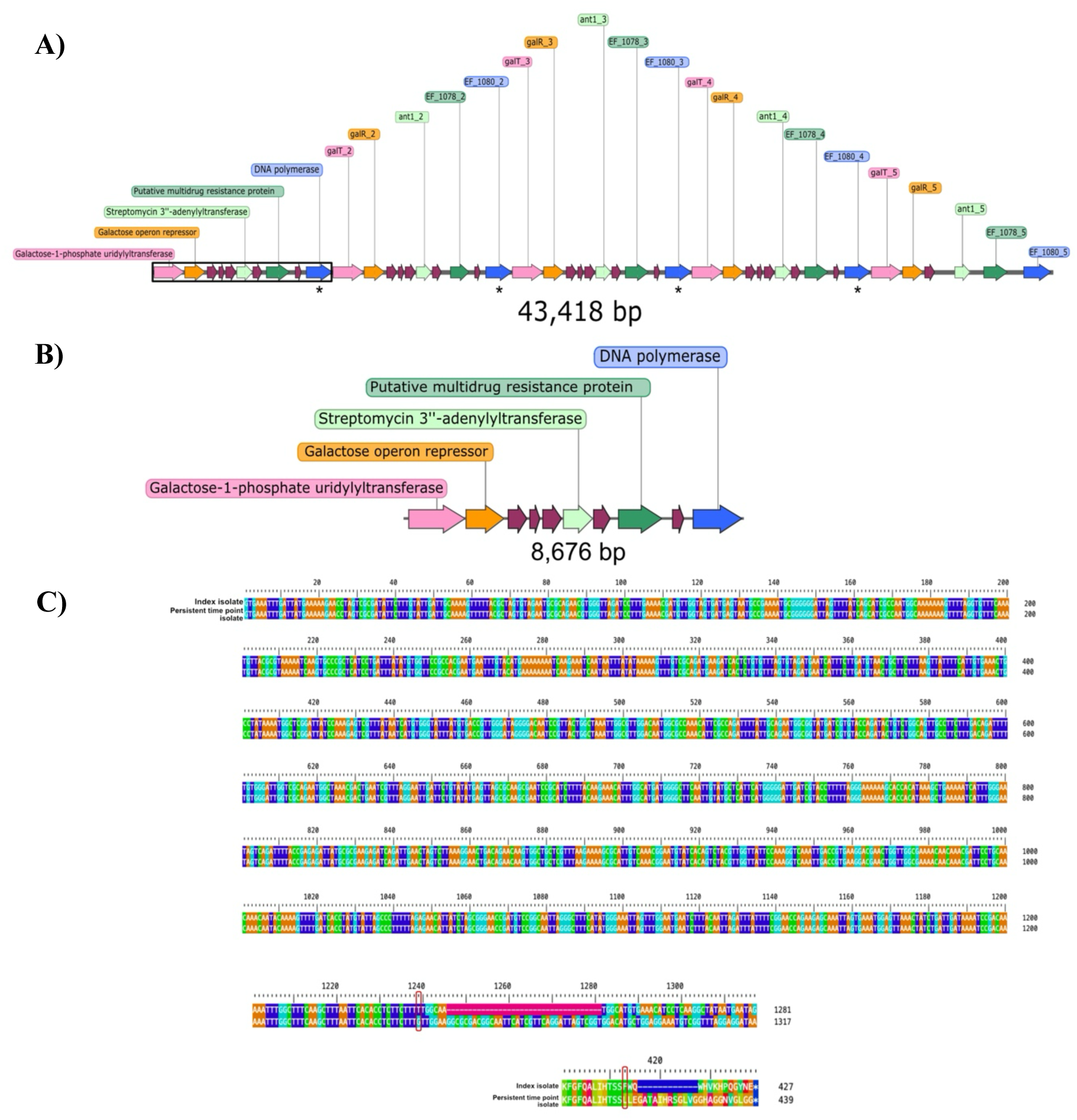
Tandem duplication of gene cluster in *E. faecalis* persistent infection pair. A) Gene map of the five-fold tandem duplication of a gene set in the index *E. faecalis* bloodstream isolate. Dark purple genes indicate hypothetical proteins. An asterisk under the gene indicates a truncation of the gene sequence. B) Gene map of the single-copy gene cluster in the persistent timepoint isolate from the same patient. C) Nucleotide alignment of the DNA polymerase gene in the index and persistent isolate demonstrating the frameshift mutation in the nucleotide sequence at position 1239, resulting in mutation L413F and an early stop codon and truncated protein.

**Figure S9.**
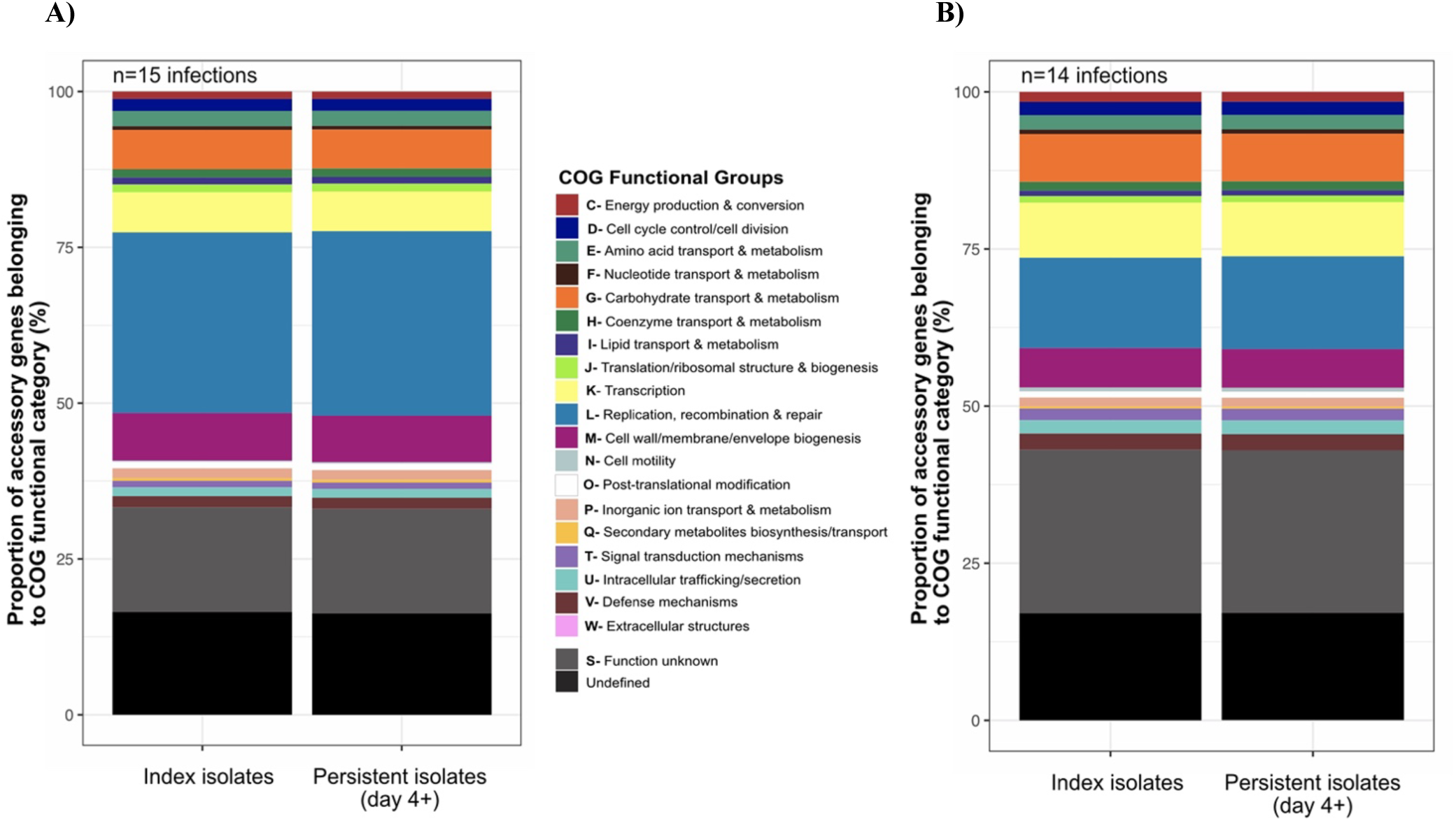
Functional characterization of accessory genes present in isolates causing persistent infections. A) *E. faecium;* B) *E. faecalis.* Stacked bar plots indicating the average proportion of the functional classes associated with accessory genes in persistent time point isolates relative to their respective index isolate.

**Figure S10.**
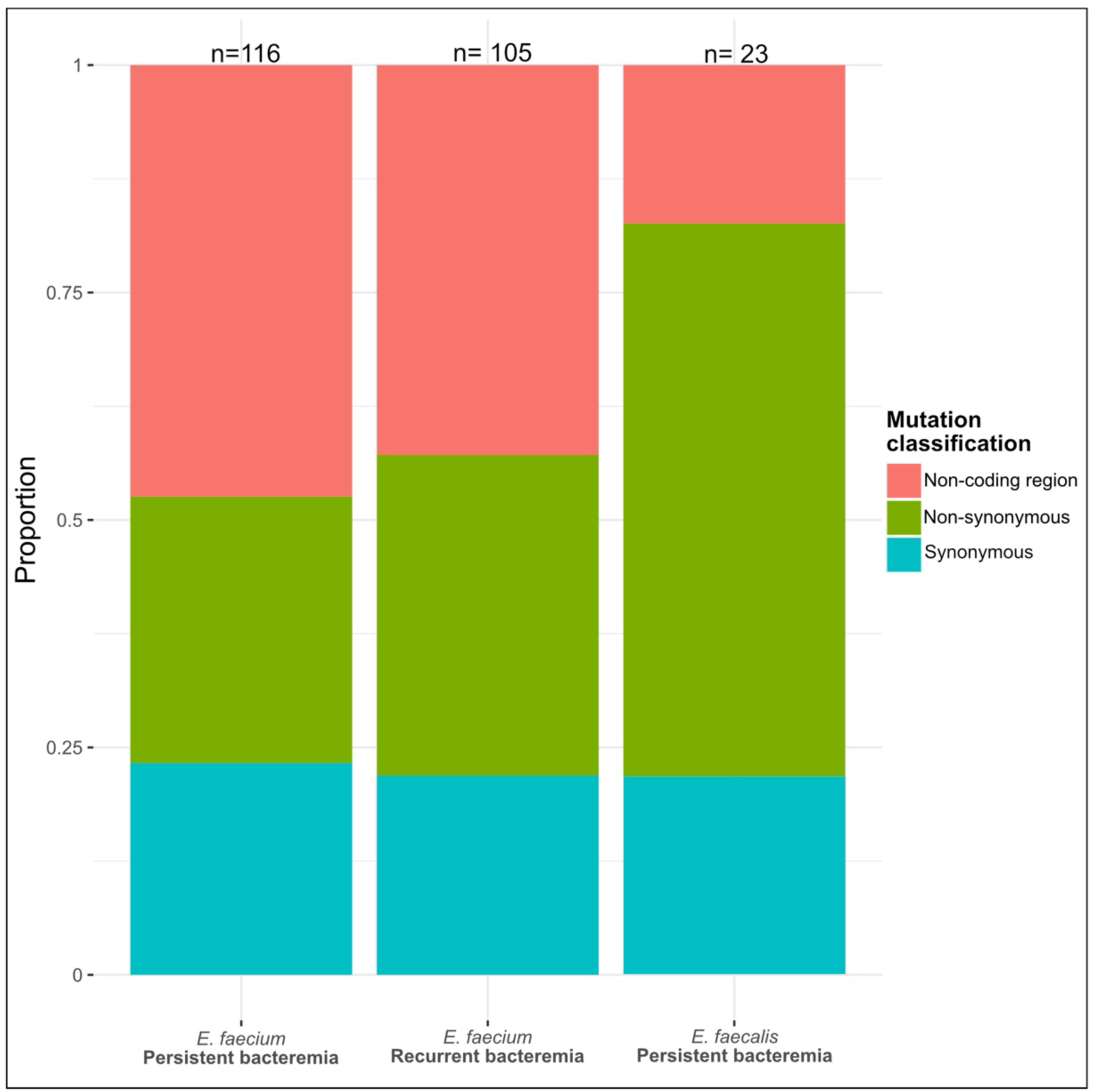
Distribution of mutation types according to species and type of bacteremia. Stacked bar plots indicating the proportion of different mutation types in bacteremia episode groups. The numbers above each bar plot indicate the total number of mutations identified in the respective group. As there were no *E. faecalis* recurrent bacteremia isolates available for sequencing, this group is absent from the figure.

**Figure S11.**
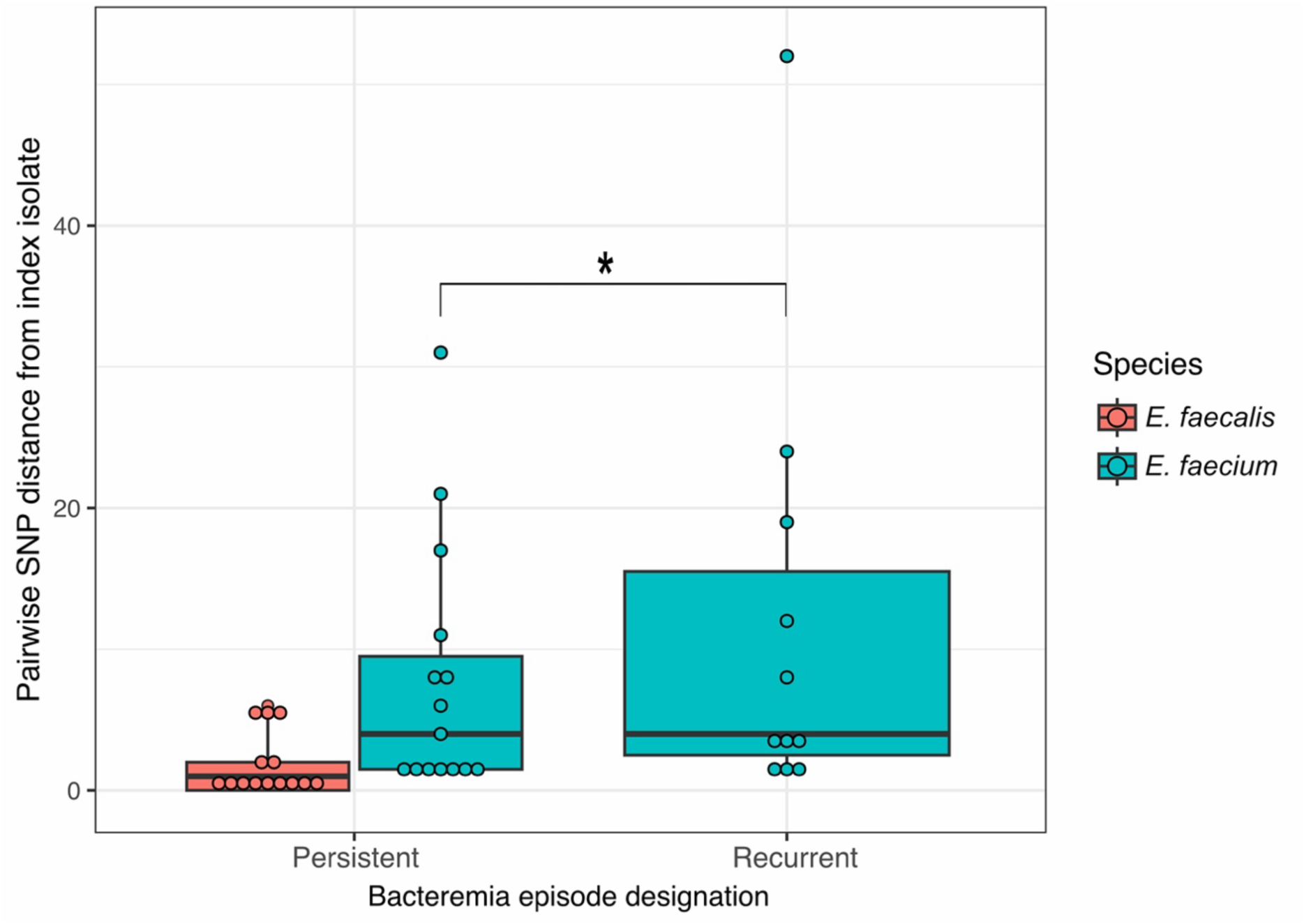
Distribution of SNPs in recurrent and persistent bacteremia isolates relative to the index isolate. Box-and-whisker plots denoting the pairwise SNP distance distributions for each bacteremia group and species. An Asterisk indicates a Wilcoxon rank-sum p-value of <0.05.

